# Inhibitors of gut bacterial L-dopa decarboxylation with reduced susceptibility to host metabolism

**DOI:** 10.64898/2026.04.08.717077

**Authors:** Rohan Narayan, Chi “Chip” Le, Jai K. Khurana, Vincent Nieto, Christine A. Olson, Peter J. Turnbaugh, Emily P. Balskus

## Abstract

Microorganisms in the human gut influence the efficacy and metabolism of host-targeted small molecule therapeutics, including the frontline Parkinson’s disease drug levodopa (L-dopa). Previous work has identified a mechanism-based inhibitor of gut bacterial decarboxylases that degrade L-dopa, α-fluoromethyltyrosine (AFMT). However, early experiments with AFMT in rodent models suggested undesirable *in vivo* metabolism by host tyrosine hydroxylase, producing α-fluoromethyldopa, a metabolite likely to worsen Parkinson’s phenotypes and prevent application as an L-dopa co-treatment. Here, we demonstrate oxidation of AFMT to α-fluoromethyldopa *in vitro* by recombinant human tyrosine hydroxylase. We then develop AFMT analogs that retain activity against bacterial decarboxylases but have reduced susceptibility to host hydroxylation. Suitable arenes for inhibitor design were identified using assays with commercially available noncanonical amino acids, which revealed aryl difluorination as a promising modification. Difluoroaryl AFMT derivatives are less prone to degradation by tyrosine hydroxylase *in vitro* yet still inhibit L-dopa metabolism by bacterial decarboxylases. This work exemplifies how substrate reactivity can streamline design of mechanism-based enzyme inhibitors, as well as how constraints posed by the host can be incorporated during development of microbiome-targeted therapeutics. The compounds reported here are promising starting points for future studies in animal models and further exploration of gut bacterial effects on L-dopa treatment efficacy.

## Introduction

Catecholamines are endogenous neurochemicals that play diverse regulatory and signaling roles in the human body. Their biosynthesis begins with hydroxylation of the aryl ring of tyrosine by a pterin-dependent tyrosine hydroxylase (TH) enzyme, producing the catecholic amino acid 3,4-dihydroxy-L-phenylalanine (levodopa or L-dopa, **1** in Fig. 1A)^1,2^. L-Dopa is then decarboxylated by the pyridoxal 5’-phosphate (PLP)-dependent enzyme aromatic amino acid decarboxylase (AADC, also known as dopa decarboxylase), forming dopamine, an important neurotransmitter in the central nervous system and a precursor to other catecholamines (Fig. 1A–1C).

**Figure 1.**
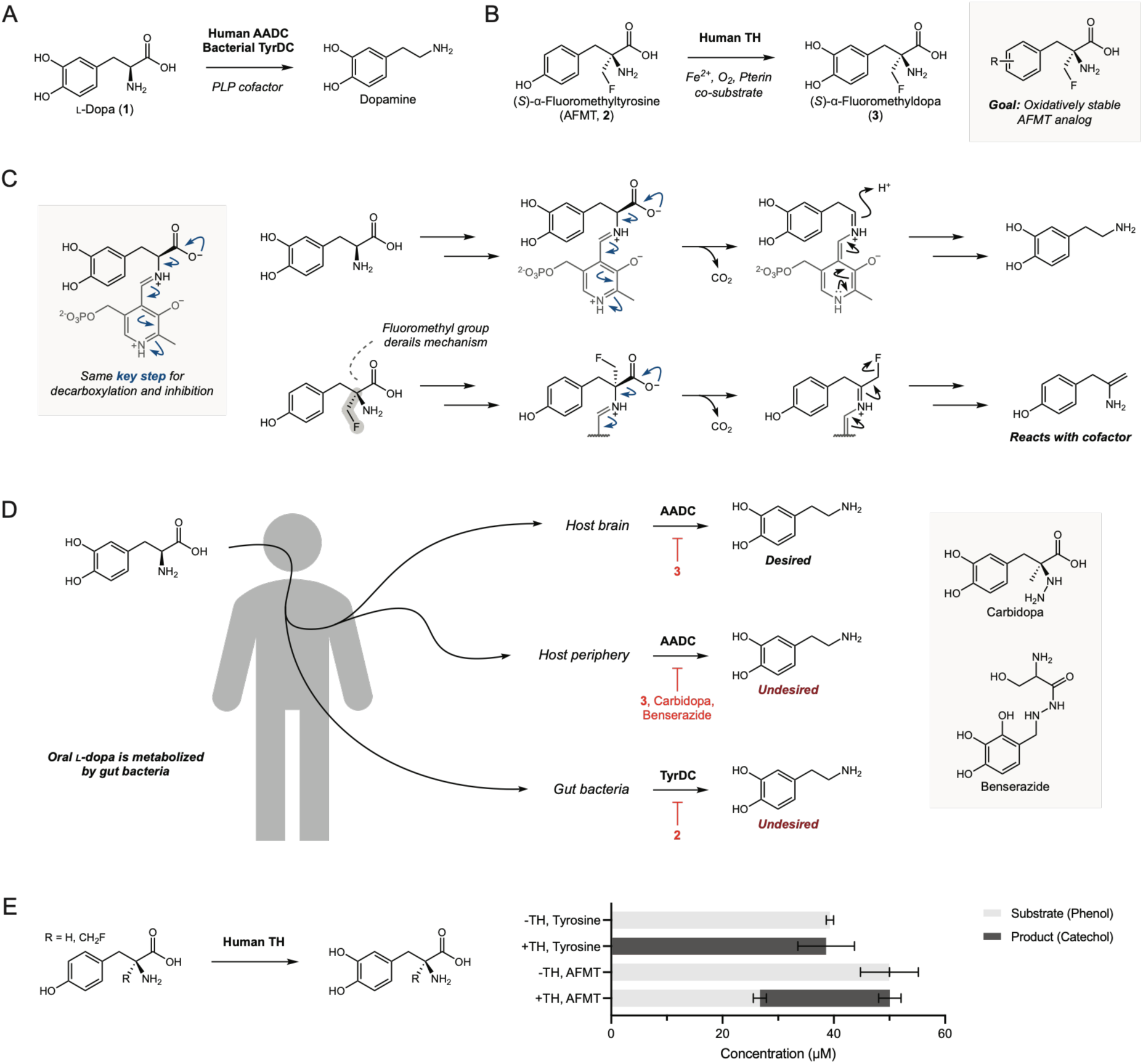
A metabolite of the gut microbial TyrDC inhibitor AFMT inhibits L-dopa decarboxylation in the mammalian brain. (*A*) The Parkinson’s disease drug L-dopa can be decarboxylated to dopamine by human AADC or gut bacterial TyrDC enzymes. (*B*) The α-fluoromethyl amino acid AFMT is thought to be hydroxylated by mammalian TH enzymes, producing the brain-penetrant compound **3**. AFMT and **3** are known inhibitors of *E. faecalis* TyrDC activity and mammalian AADC activity, respectively. This work seeks to identify AFMT analogs with reduced susceptibility to hydroxylation. (*C*) L-Dopa decarboxylation by PLP-dependent enzymes begins with iminium formation connecting substrate and cofactor. Decarboxylation is enabled by the electron-deficient pyridinium ring, after which the resultant quinonoid intermediate is protonated at the α-carbon. Inhibition by AFMT is thought to proceed through an analogous key decarboxylation step, with fluoride loss from the quinonoid intermediate instead producing a nucleophilic enamine product. (*D*) L-Dopa is administered orally to manage Parkinson’s disease, with dopamine production in the brain being the desired outcome. Carbidopa, benserazide, and **3** are known inhibitors of mammalian AADC activity, and the former two are co-administered with L-dopa to prevent AADC-mediated decarboxylation outside the brain. Gut bacteria, such as *E. faecalis*, can decarboxylate L-dopa prior to absorption through the action of TyrDC enzymes. Co-administration of AFMT with L-dopa could enhance frontline Parkinson’s treatment by inhibiting bacterial decarboxylase activity. (*E*) *In vitro* assay with human TH showing conversion of tyrosine to L-dopa and AFMT to **3**. 0.5 µM human TH was incubated with 50 µM phenol substrate (tyrosine or AFMT) and appropriate cofactors for two hours at 37 °C in pH 7 HEPES buffer. Final concentrations of phenol substrates and catechol products were determined using ultra-performance liquid chromatography-tandem mass spectrometry (UPLC–MS/MS) and are shown as stacked bars where the length of each segment corresponds to analyte concentration. Error bars show mean ± standard deviation of triplicate assays run in the same plate, and results are representative of two experiments conducted on different days.

A loss of dopamine-producing neurons in the brain’s substantia nigra drives the motor symptoms associated with Parkinson’s disease, a neurological disorder projected to affect more than 1.2 million individuals in the United States by 2030^3,4^. Because dopamine itself cannot cross the blood-brain barrier, these symptoms are typically treated through oral administration of the dopamine precursor L-dopa^2,5,6^. L-Dopa is absorbed in the intestines and transported into the brain where it is decarboxylated by AADC to restore the depleted dopamine levels (Fig. 1D). AADC is also expressed outside the central nervous system, so L-dopa is usually coadministered with carbidopa or benserazide, AADC inhibitors that prevent drug consumption in the periphery but cannot cross the blood-brain barrier^2,3,7,8^. AADC inhibitors have greatly enhanced the efficacy of oral L-dopa treatment, and these drug combinations are currently the frontline standard for Parkinson’s disease management. However, even with carbidopa co-administration as little as 60% of ingested L-dopa may enter the bloodstream unmodified, with an even smaller fraction (∼10%) reaching the brain^3,9^. This residual peripheral metabolism is thought to reduce drug efficacy, necessitate higher dosing, and contribute to undesirable GI side effects.

Our group and others previously identified human gut microbes that decarboxylate L-dopa and may contribute to undesirable drug metabolism^10,11^. An intracellular, PLP-dependent tyrosine decarboxylase (TyrDC) enzyme encoded by gut enterococci was shown to decarboxylate L-dopa *in vitro* and in bacterial culture. Unlike host AADC, this enzyme was not inhibited by carbidopa. *Ex vivo* L-dopa metabolism in fecal samples from Parkinson’s patients showed high interpersonal variability, consistent with an effect driven by microbiome composition. Importantly, fecal *tyrDC* gene abundance associates with differences in *ex vivo* L-dopa metabolism, as well as with clinical variables such as increased disease duration, increased daily drug dose, lower *in vivo* drug exposure, and poorer responsiveness to treatment^10–15^. Recent reports have also linked gut bacterial L-dopa decarboxylation to decreased drug efficacy in a chemically-induced animal model of Parkinson’s disease^16,17^. Together, these findings suggest gut bacterial decarboxylation could be a relevant route of peripheral L-dopa metabolism in patients and point to an opportunity for improving treatment by inhibiting TyrDC.

We previously identified (*S*)-α-fluoromethyltyrosine (AFMT, **2**) as an effective mechanism-based inhibitor of TyrDC activity in *Enterococcus faecalis* culture, human fecal samples, and a gnotobiotic animal model. The proposed mechanism for inhibition parallels standard catalysis through the decarboxylation step, however instead of protonation at the α-carbon, AFMT can lose an equivalent of fluoride to produce an enamine which inactivates the cofactor^10^. AFMT shows little to no inhibitory activity against human AADC, yet earlier literature indicates its catecholic analog (α-fluoromethyldopa, **3**) is a potent, brain-penetrant AADC inhibitor^18–20^. Prior studies suggest that AFMT can be hydroxylated to compound **3** by mammalian TH, mirroring the biosynthesis of L-dopa from tyrosine. Gavage of an AFMT methyl ester prodrug leads to slow inhibition of AADC activity in the rat brain, presumably through production of **3**, an effect that can be altered by known modulators of TH activity^21^. The same report notes that hydroxylation of AFMT was shown *in vitro* using partially purified TH from porcine adrenal glands. These observations have since been utilized to design AFMT derivatives as radiotracers for TH activity^22–24^. Because brain AADC activity is critical for L-dopa treatment, *in vivo* production of **3** presents a potential liability for therapeutic use of AFMT.

Here, we aim to address this problem by developing AFMT analogs that are less susceptible to oxidation by TH. We first demonstrate AFMT hydroxylation by human TH *in vitro*. We then use TyrDC’s activity toward noncanonical amino acids in bacterial culture and TH’s activity *in vitro* to design difluoroaryl AFMT analogs that are less susceptible to TH metabolism. We observe differences in oxidative stability and TyrDC inhibition potency for the different analogs, and we demonstrate that these compounds inhibit L-dopa metabolism by multiple gut enterococci and human fecal samples. This work sets the stage for future investigation of gut bacterial TyrDC inhibitors as therapeutic candidates to enhance L-dopa therapy.

## Results

### 3,5-Difluorotyrosine is accepted by TyrDC but is a poor substrate for TH

There is already strong evidence in the literature for hydroxylation of AFMT by mammalian TH, but we were unable to find a clear demonstration of this metabolism by the human enzyme^21,23,24^. We therefore began by confirming that human TH hydroxylates AFMT to produce compound **3** *in vitro*. The human enzyme was heterologously expressed in *Escherichia coli* as a maltose-binding protein fusion^25^. The fusion partner was removed with TEV protease, after which the purified TH was assayed for AFMT hydroxylation using ultra-performance liquid chromatography-tandem mass spectrometry (UPLC–MS/MS). AFMT consumption was observed upon incubation with TH, and production of **3** was confirmed through comparison with a synthetic standard (Fig. 1E). Hydroxylation of the α-fluoromethyl inhibitor was slower than the analogous reaction of tyrosine, with close to half the substrate converted after two hours under our reaction conditions, consistent with work by Jung et al. using partially purified porcine enzyme^21^. Our *in vitro* result supports the precedent that mammalian TH can produce a potent AADC inhibitor from AFMT.

We reasoned that modifying electronic properties of the aromatic ring of AFMT and/or blocking potential reactive sites could provide an inhibitor which maintains efficacy against *E. faecalis* TyrDC but is less susceptible to hydroxylation by human TH. Since AFMT derivatives are resource-intensive to make and purify, we instead evaluated commercially available noncanonical amino acids as candidate substrates for TyrDC (Fig. 2A). Decarboxylation is the first step in enzyme inactivation by α-fluoromethyl amino acids, so we expected that α-fluoromethyl analogs of amino acids which are robustly metabolized by *E. faecalis* TyrDC would have inhibitory activity against this enzyme. This substrate-inhibitor analogy is consistent with logic employed in early studies of α-fluoromethyl decarboxylase inhibitors^18^. We also expected the susceptibility of an aromatic amino acid to hydroxylation by TH would mirror that of the corresponding α-fluoromethyl amino acid, allowing us to evaluate relative oxidative stability without needing to synthesize each inhibitor. Development of AFMT analogs therefore proceeded in stages: 1) identifying amino acids that are decarboxylated in *E. faecalis* culture, 2) checking for metabolism of promising substrates by TH, and 3) incorporating select aryl substituents into a focused set of candidate inhibitors (Fig. 2B). We hypothesized that amino acids which are metabolized by *E. faecalis* TyrDC but not hydroxylated by human TH would be ideal starting points for inhibitor design.

**Figure 2.**
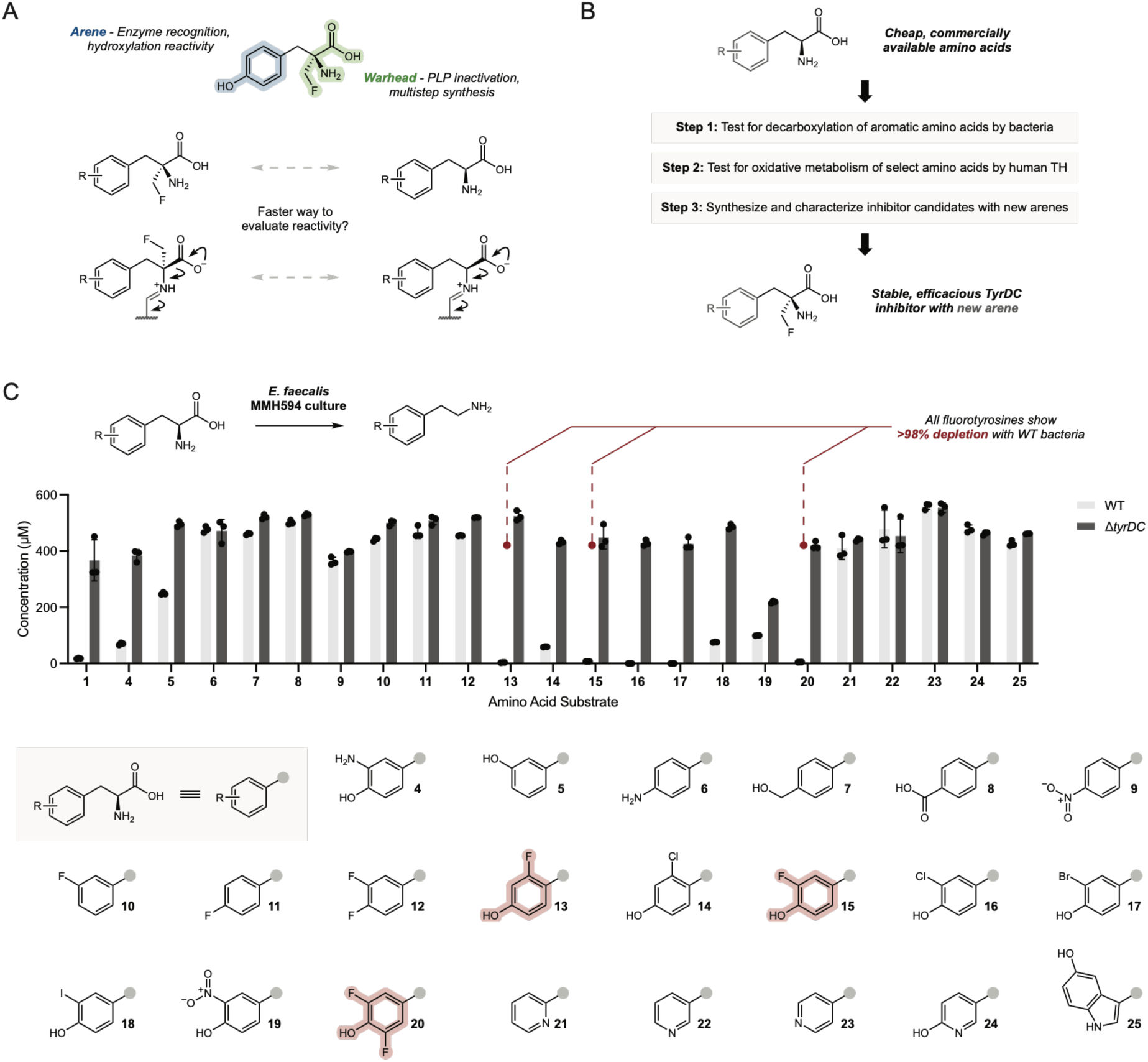
Fluoroaryl tyrosine derivatives exhibit TyrDC-dependent depletion in *E. faecalis* culture. (*A*) AFMT can be thought of as two fragments: 1) an aryl portion putatively responsible for enzyme recognition and hydroxylative reactivity, and 2) a PLP-reactive α-fluoromethyl amino acid warhead that confers inhibitory activity. The inhibition mechanism initiates analogously to amino acid decarboxylation, suggesting that decarboxylation efficiency of aromatic amino acids would parallel inhibitory activity of the analogous α-fluoromethyl compounds. (*B*) Considering the substrate-inhibitor analogy outlined in panel *A*, we used decarboxylation assays with noncanonical amino acids to identify promising scaffolds for inhibitor development. We evaluated the hydroxylative reactivity of multiple fluorotyrosines using *in vitro* assays with human TH, after which a focused set of α-fluoromethyl amino acids was synthesized and evaluated for TyrDC inhibition. (*C*) Testing noncanonical amino acids for TyrDC-mediated metabolism in bacterial culture. *E. faecalis* MMH594 WT or Δ*tyrDC* mutant strains were grown for 18 hours under anaerobic conditions with 500 µM of each amino acid substrate shown. The final amino acid concentration was determined using UPLC–MS/MS, and error bars show mean ± standard deviation of triplicate cultures grown in the same plate.

To begin this study, *E. faecalis* MMH594 was grown in the presence of 22 different noncanonical aromatic amino acids (500 µM) for 18 hours and substrate concentrations post-incubation were determined using UPLC–MS/MS (Fig. 2C). Testing for decarboxylation in bacterial culture instead of with purified enzyme allowed us to identify not only which substituents are tolerated by the decarboxylase, but also those which are accepted by any relevant bacterial transporters. This is important for inhibitor design, since TyrDC is intracellular and transport limitations can be formidable in bacteria. We found that several electron-deficient tyrosine derivatives were consumed after incubation with *E. faecalis* (**13**–**20**), and in particular all three fluorotyrosines exhibited even lower final concentrations than L-dopa (**13**, **15**, **20**). To increase our confidence that the TyrDC enzyme was responsible for the observed depletion, an *E. faecalis* MMH594 Δ*tyrDC* strain was grown with the same panel of amino acid substrates. Comparing these results to those from the wild type (WT) strain shows that TyrDC is required for depletion of most substrates, with compound **19** being an obvious exception. We hypothesized that reduction of the nitro group of **19** could explain this finding since *E. faecalis* is known to encode functional nitroreductases^26^. Given that compound **9** also features a nitroarene, we investigated and confirmed production of the cognate anilines after incubation of *E. faecalis* with either nitro-containing substrate (Figure S5).

As mentioned above, all three fluorotyrosine substrates were efficiently metabolized during the decarboxylation assays in bacterial culture. This drew our attention, given that fluorination is a precedented strategy to disfavor oxidative metabolism of arenes, and we wondered if incorporation of fluorophenol motifs could slow TH-mediated hydroxylation of AFMT-like compounds^27,28^. We therefore progressed the three fluorotyrosine substrates to assays with purified human TH (Fig. 3A). 2-Fluorotyrosine (**13**) was completely depleted after one hour under the assay conditions (as was the tyrosine control), and 3-fluorotyrosine (**15**) showed more than 80% consumption at the same timepoint. In contrast, 3,5-difluorotyrosine (**20**) showed less than 20% consumption after one hour, suggesting difluorinated arenes could furnish inhibitors with decreased hydroxylation susceptibility. These results are consistent with those reported by Hillas and Fitzpatrick using the catalytic domain of a rodent TH, with the addition of a difluoroaryl substrate^29^. Hydroxylation of compound **15** is also consistent with work by DeJesus et al., who showed that an AFMT analog with a radiolabeled 3-fluoro arene is still converted to a viable AADC inhibitor by TH in rat brain homogenates^22^.

**Figure 3.**
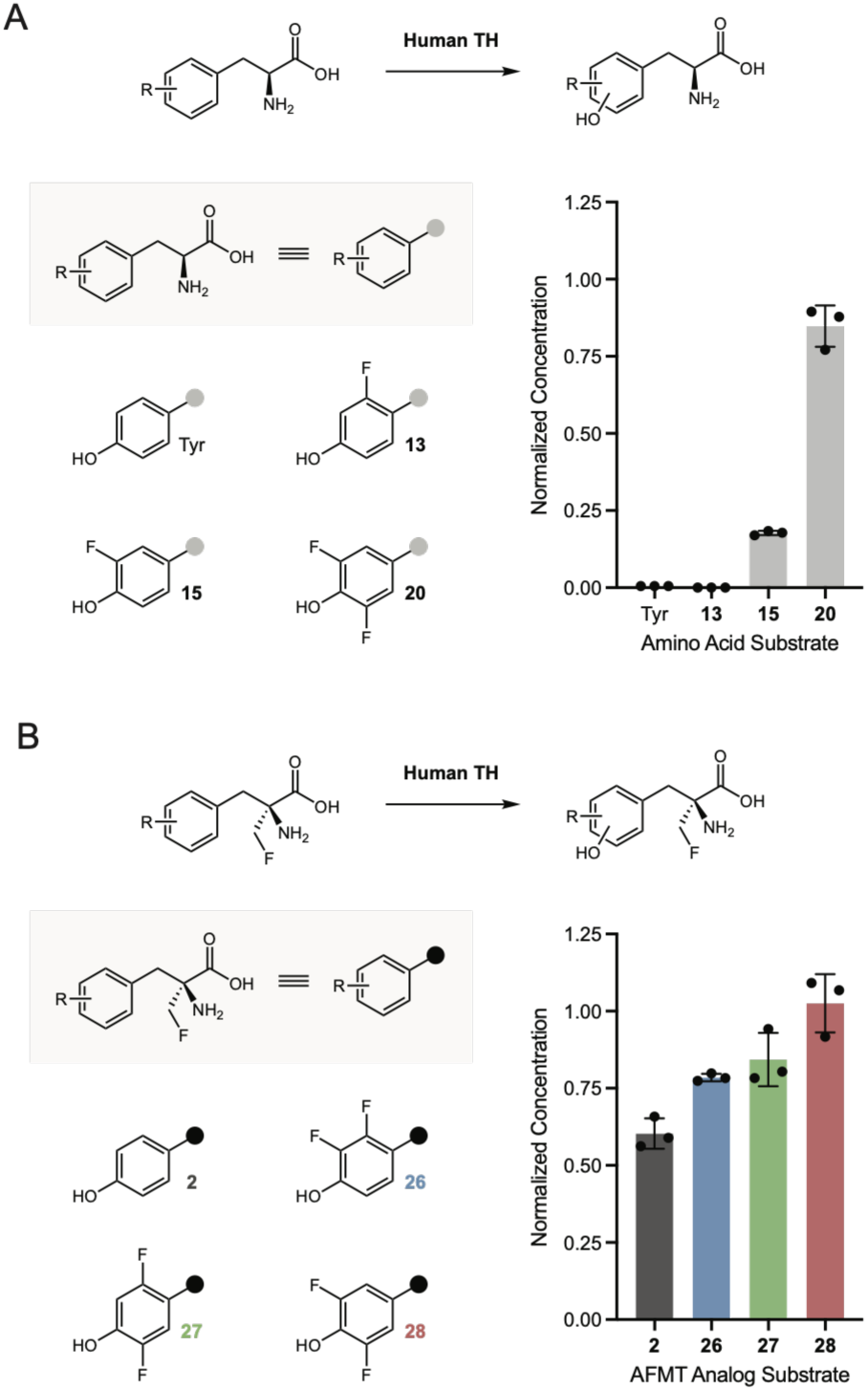
Difluoroaryl amino acid derivatives show reduced susceptibility to metabolism by TH. (*A*) Testing fluorotyrosines for hydroxylation by TH. Amino acids (50 µM) were incubated with 0.5 µM purified human TH and appropriate cofactors for one hour at 37 °C in pH 7 HEPES buffer. Substrate remaining was quantified using UPLC–MS/MS, and the values shown are normalized to those observed in no-enzyme controls. (*B*) Testing AFMT and difluoroaryl analogs for hydroxylation by TH. Assay conditions and normalization are the same as described in *A*, except with a two-hour incubation and altered quench conditions. Error bars for both *A* and *B* show mean ± standard deviation of triplicate assays run in the same plate, and results in both panels are representative of two experiments conducted on different days.

### Difluoroaryl AFMT analogs are poor substrates for TH

We elected to use the difluorotyrosine scaffold as a starting point for designing TyrDC inhibitors with reduced susceptibility to hydroxylation by TH. Fluorophenols exhibit stark differences in p*K*_a_ and redox behavior depending on the number and position of fluoride substituents, an insight that has been applied to inhibitor design and protein biochemistry. Additionally, varying the site of aryl fluoride substitution has been shown to dramatically alter potency, selectivity, and pharmacokinetic properties for drug candidates^27,28,30–32^. We therefore chose to synthesize and evaluate a panel of three AFMT analogs with different difluorination patterns on the aromatic ring: 2,3-difluoroaryl analog **26**, 2,5-difluoroaryl analog **27**, and 3,5-difluoroaryl analog **28**.

Before inhibitor characterization, we sought to verify that aryl difluorination would indeed reduce hydroxylation of the synthesized AFMT analogs by TH (Fig. 3B). We compared depletion of compounds **26**–**28** upon incubation with TH and detected all three difluoroaryl inhibitors at higher normalized concentrations than AFMT after two hours. Compound **28** appeared unmetabolized compared to a no-enzyme control at this timepoint, while **26** and **27** showed ∼80% substrate remaining compared to ∼60% for AFMT. This result suggests that difluoroaryl AFMT analogs may be less susceptible to metabolism by TH upon *in vivo* administration, potentially enabling therapeutic application without inhibition of brain AADC activity.

### Difluoroaryl AFMT analogs are potent TyrDC inhibitors

We next evaluated the difluoroaryl AFMT derivatives for inhibition of TyrDC in bacterial culture and *in vitro*. *E. faecalis* MMH594 was grown in the presence of 500 µM L-dopa and varying concentrations of compounds **26**–**28**, after which remaining L-dopa was quantified using UPLC– MS/MS (Fig. 4A). While analogs **26** and **27** displayed comparable efficacy to AFMT (EC_50_ ∼1 µM), compound **28** showed a striking loss of potency. We were curious whether this decrease was attributable to less potent TyrDC inhibition by compound **28** or to poorer uptake by bacterial cells. We therefore heterologously expressed and purified the *E. faecalis* MMH594 TyrDC and used the purified protein for *in vitro* assays with compounds **26**–**28** (Fig. 4B). Compound **28** was also a less potent inhibitor of L-dopa decarboxylation when tested against purified protein, suggesting reduced activity toward TyrDC is responsible for the decreased potency observed in culture.

**Figure 4.**
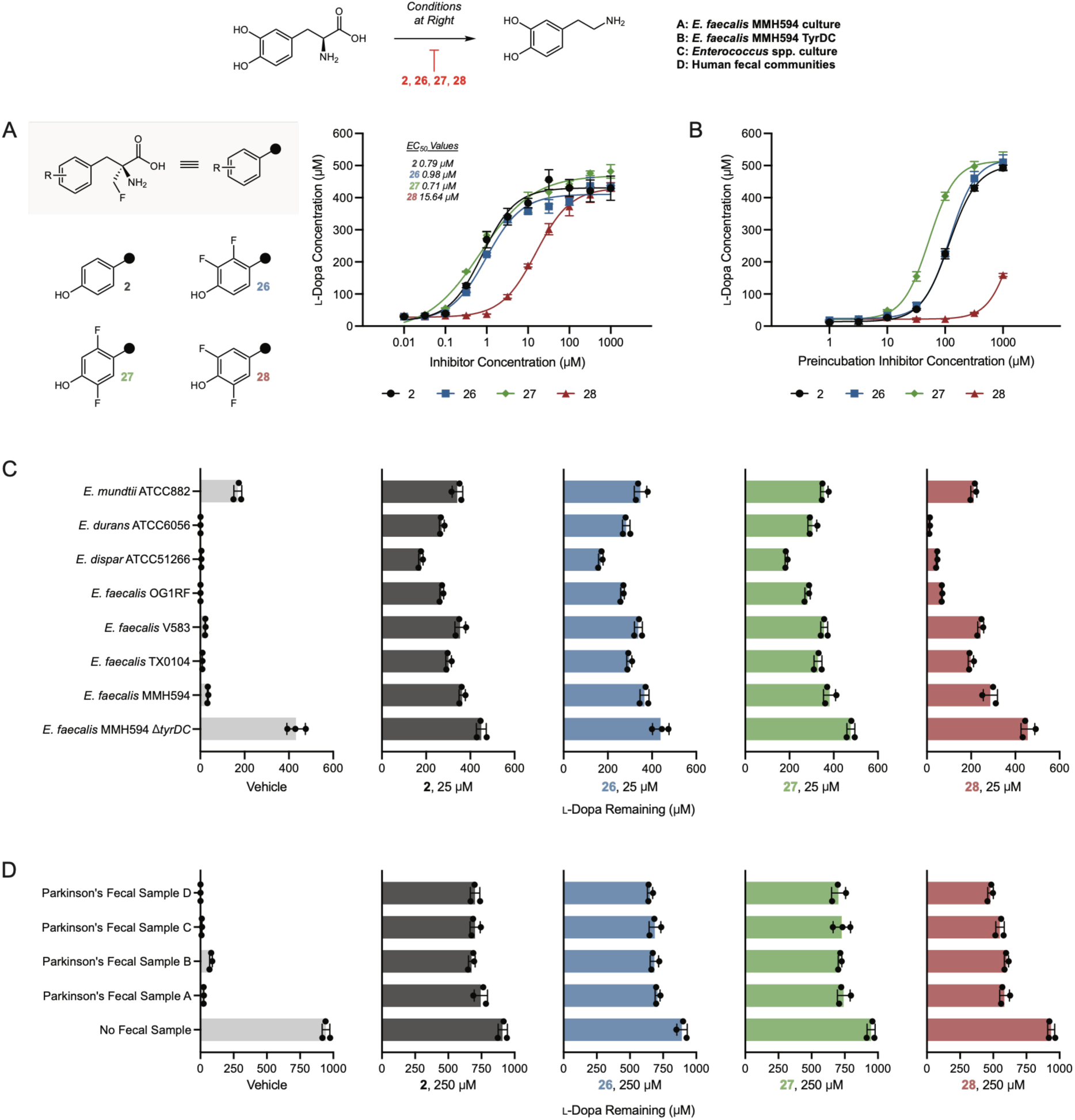
Difluoroaryl AFMT analogs inhibit TyrDC. (*A*) Inhibition of L-dopa decarboxylation by AFMT and difluoroaryl analogs in bacterial culture. *E. faecalis* MMH594 was grown for 18 hours with 500 µM L-dopa and varying inhibitor concentrations. L-Dopa remaining post-incubation was quantified using UPLC–MS/MS. (*B*) Inhibition of L-dopa decarboxylation by AFMT and difluoroaryl analogs *in vitro* with purified *E. faecalis* TyrDC. 1 µM purified TyrDC was preincubated for 20 minutes with 200 µM PLP and varying inhibitor concentrations at room temperature in pH 7 HEPES buffer. The preincubations were then diluted 10X in a solution of substrate to give assay concentrations of 500 µM L-dopa, 0.1 µM TyrDC, 20 µM PLP, and varying inhibitor concentrations in pH 5.5 sodium acetate buffer. Remaining substrate after 20 minutes of reaction was quantified using UPLC–MS/MS. Note that inhibitor concentrations used for preincubation are plotted. (*C*) Inhibition of L-dopa decarboxylation by AFMT and difluoroaryl analogs in assays with a panel of *Enterococcus* strains. Each strain was grown for 18 hours in the presence of 500 µM L-dopa and 25 µM inhibitor, after which remaining L-dopa was quantified using UPLC–MS/MS. (*D*) Inhibition of L-dopa decarboxylation by AFMT and difluoroaryl analogs in complex communities derived from Parkinson’s patient fecal samples. Each community was grown for 40 hours in the presence of 1000 µM L-dopa and 250 µM inhibitor, after which remaining L-dopa was quantified using UPLC–MS/MS. All four samples were provided by distinct donors. For all panels, error bars show mean ± standard deviation of triplicate assays run in the same plate, and for panels *A*–*C* results are representative of two experiments conducted on different days. All experiments involving bacterial growth were conducted under anaerobic conditions.

We lastly sought to address two important uncertainties when considering co-administration of inhibitors **26**–**28** with L-dopa: 1) possible AADC inhibition by the unmetabolized compounds and 2) efficacy against TyrDC enzymes encoded by other human gut bacteria. We reasoned that host AADC was a plausible off-target for mechanism-based TyrDC inhibitors, given the similar reactions performed by these two enzymes. Although AADC inhibition outside the central nervous system is potentially favorable for L-dopa therapy, inhibition in the brain is very undesirable, and we cannot guarantee our compounds would be peripherally restricted without a full pharmacokinetic study. Additionally, though AFMT itself is not a competent AADC inhibitor, fluorophenol analogs of carbidopa have been proposed for targeting the host enzyme, further motivating experiments testing our difluoroaryl AFMT analogs against AADC^33^. We therefore expressed and purified the human AADC from *E. coli* and tested for inhibition by AFMT and the difluoroaryl analogs (Figure S8). We did not observe AADC inhibition with any of these compounds under our assay conditions, in stark contrast to the carbidopa positive control.

To investigate inhibitor efficacy against additional bacteria, we grew a panel of TyrDC-encoding enterococci in the presence of 500 µM L-dopa with or without 25 µM inhibitor, then quantified L-dopa remaining (Fig. 4C). Compounds **26** and **27** demonstrated similar efficacy to AFMT at the tested concentration for all organisms in the panel. Compound **28** again showed lower efficacy than the other inhibitors in these experiments. Post-incubation OD_600_ values were similar with or without inhibitor treatment, suggesting inhibition of TyrDC rather than compound toxicity is responsible for the increased L-dopa remaining (Figure S9). Finally, to explore inhibitor efficacy in complex human gut microbiome samples, we tested compounds **26**–**28** in *ex vivo* assays using fecal samples from Parkinson’s patients (Fig. 4D). In this case the L-dopa substrate was provided at 1000 µM and inhibitors were tested at 250 µM. We observed similar trends to the single-strain experiments, with compounds **26** and **27** displaying comparable efficacy to AFMT and compound **28** exhibiting weaker inhibition. Taken together, these inhibition assays suggest difluoroaryl AFMT analogs can prevent L-dopa decarboxylation by gut bacteria without unwanted inhibition of host AADC.

## Discussion

Gut bacterial TyrDC enzymes are known to decarboxylate the frontline Parkinson’s disease drug L-dopa, and multiple publications have observed associations between *tyrDC* gene abundance in patient stool and clinical variables such as treatment response, drug bioavailability, and disease duration^10–15^. This precedent suggests a connection between TyrDC activity and clinical L-dopa efficacy, though further studies are warranted to better characterize the scope and importance of this metabolism in a clinical setting^34^. Other gut bacterial enzymes have been reported to metabolize L-dopa, and improvements in drug pharmacokinetics or treatment response were transferrable through fecal microbiome transplant in a mouse model, indicating that gut bacteria present an important target for improved L-dopa therapy^17,35–37^. Methods are now being proposed to manipulate TyrDC activity in the gut through selective killing of enterococci or small molecule inhibitor development^16,38–40^. Reunanen et al. recently highlighted a chemically distinct compound class with activity against both TyrDC and AADC *in vitro*, and inhibition of both enzymes using a single molecule could present an attractive alternative to inhibitor co-administration (ex. AFMT dosing alongside carbidopa)^40^. However, the same report notes that the disclosed compounds display concerning toxicity towards human and bacterial cells, the inhibition of TyrDC appeared less potent than with AFMT, and inhibitor characterization in bacterial culture was limited to one strain of *E. faecalis* under aerobic conditions.

Here, we add to this exciting space by identifying new gut bacterial TyrDC inhibitors as candidates for co-administration with L-dopa. AFMT analogs with substitution on the aromatic ring were envisioned to avoid hydroxylation by host TH and minimize unwanted inhibition of AADC activity in the brain. By assaying a panel of commercially available noncanonical amino acids, we quickly identified a difluorotyrosine substrate of *E. faecalis* TyrDC that is poorly metabolized by the mammalian hydroxylase. Multiple difluoroaryl AFMT analogs were synthesized and confirmed to be poorer substrates for TH, with the 3,5-difluoroaryl inhibitor (**28**) displaying the greatest decrease in oxidative susceptibility. However, compound **28** was more than 15-fold less potent for inhibition of L-dopa decarboxylation by *E. faecalis* culture than its regioisomers or AFMT. This potency difference likely arises from decreased activity toward TyrDC itself, since efficacy against purified enzyme showed a similar trend to that observed in culture. Notably, the 2,3-difluoroaryl and 2,5-difluoroaryl AFMT analogs (**26** and **27**) showed similar potency to AFMT in bacterial culture and against purified TyrDC, though both displayed increased stability to TH.

Noncanonical amino acids were used for initial assays with bacterial culture and purified enzyme, allowing us to efficiently identify and rule out scaffolds unlikely to produce potent or metabolically stable inhibitors before pursuing synthesis. α-Fluoromethyl amino acids require bespoke synthesis and chiral separation, so initial synthesis of all candidate inhibitors would have been time-and resource-intensive. Our alternative approach was enabled by two key observations: 1) since α-fluoromethyl amino acids are mechanism-based inhibitors, their inhibitory activity should parallel decarboxylation efficiency of the unmodified amino acids, and 2) a wide variety of noncanonical aromatic amino acids are commercially available. This strategy greatly streamlined medicinal chemistry efforts by focusing synthetic resources on a narrow set of target compounds.

Difluorophenol regioisomers have been explored in several medicinal chemistry campaigns to date, and those studies have shown that changing the sites of fluorination can have remarkable impact on compound efficacy^41–47^. The differences we observe in potency and metabolic stability among our TyrDC inhibitors are consistent with this precedented sensitivity to fluorination pattern. Based on prior application of fluorotyrosine regioisomers in mechanistic enzymology, we expect differences in phenol acidity, arene electronics, and redox behavior among our AFMT analogs^48,49^. Compound **28** features the most acidic phenol, with an expected p*K*_a_ (∼6.8) roughly three pH units lower than that of AFMT (∼10) by analogy to the corresponding tyrosines^48^. Compounds **26** and **27** are expected to display phenolic p*K*_a_’s (both ∼7.6) between these two extremes. Indeed, the difluorophenol substructure of compound **28** has been invoked as a carboxylate bioisostere due in part to this enhanced acidity, highlighting the expected negative charge under physiological conditions^50^. The difluoroaryl inhibitors are also expected to have more electron-deficient aromatic ring systems than AFMT, and previous work has reported increased single-electron reduction potentials for the analogous tyrosyl radicals^49^.

Bearing these chemical properties of our TyrDC inhibitors in mind, we can hypothesize several potential mechanisms underlying the observed trends in their oxidative stability and inhibition potency. As alluded to previously, decreased *in vitro* metabolism of difluoroaryl AFMT analogs by human TH could be explained by fluorination physically blocking access to the hydroxylation site or disfavoring a catalytically productive binding pose for the substrate. Aryl difluorination could alternatively decrease binding affinity of AFMT-like substrates for TH, perhaps due in part to changes in phenolic p*K*_a_. The proposed mechanism for TH catalysis invokes a cationic intermediate formed by an electrophilic aromatic substitution event involving the substrate arene and a highly reactive ferryl oxo species (Figure S3)^1,29,51^. It could therefore be that difluorination further deactivates the aromatic ring of AFMT through electron-withdrawing effects, which would destabilize this cationic intermediate and disfavor reaction with the ferryl oxo. The improved stability of compound **28** relative to its regioisomers in experiments with TH indicates that obstruction or deactivation of both carbons *ortho* to the phenol is beneficial. Further experimentation is required to determine the exact products of TH-mediated inhibitor metabolism given the possibility for multiple sites of attack and downstream rearrangements^29,52^. In particular, the regiochemistry of oxidation for compounds **26** and **27** is of interest, since for these substrates there is the option for either C–H or C–F cleavage adjacent to the phenol.

Focusing on trends in TyrDC inhibition, only compound **28** showed a sharp drop in potency compared to AFMT, both in whole cell and purified enzyme settings. Since decarboxylation and inhibitory activity do not directly involve the aryl portion of AFMT, we hypothesize that the loss of potency is due to poorer inhibitor binding to its target enzyme. As noted above, compound **28** is also the AFMT analog with the lowest expected phenol p*K*_a_. It is therefore possible that the presence of an O–H bond, perhaps as a hydrogen bond donor, or the absence of a negative charge improves inhibitor binding to TyrDC. The relative efficacies of AFMT analogs were consistent across our panel of enterococci, supporting the notion that inhibitor recognition could be similar across the tested strains (TyrDC homolog identity is ∼60% or higher at the amino acid level). Assays with human fecal samples also showed the same pattern of inhibitor efficacy, indicating that enterococci are primary contributors to *ex vivo* L-dopa metabolism or that inhibitor recognition is similar for other relevant decarboxylase enzymes.

Though these inhibitors present an attractive starting point for therapeutic development, future work is necessary to address outstanding limitations and knowledge gaps. As mentioned earlier, additional studies could help clarify the extent of gut bacterial L-dopa metabolism and its relevance for treatment efficacy in diverse populations. Experiments with animal models have made some exciting progress in modeling the bacterial contribution to L-dopa pharmacokinetics, and in the future these models should be utilized to compare and prioritize clinical candidates for coadministration^10,11,16,17^. Animal models would also enable interrogation of the impact of aryl fluorination on the *in vivo* stability and ADME properties of our TyrDC inhibitors, including the possibility of additional metabolism^31^. The lower phenol p*K*_a_ expected for difluoroaryl AFMT analogs could result in differences in absorption and distribution, as observed by DeJesus et al. in their experiments with 3-fluoro AFMT^22^.

Overall, our work presents proof of concept for designing gut bacterial TyrDC inhibitors with reduced susceptibility to metabolism by a human enzyme. Small molecule inhibitors targeting the gut microbiome have recently become an area of interdisciplinary interest for chemical biology, medicinal chemistry, and microbiology. The field is now witnessing second-generation inhibitor studies that have improved compound potency and pharmacokinetics or demonstrated application in complex settings^53–55^. In our case, we alter a gut bacterial enzyme inhibitor to circumvent metabolism by a specific host enzyme, and to our knowledge this is the first reported instance of designing a nonlethal gut microbiome-targeted inhibitor around a host metabolic constraint. This work therefore represents important precedent in balancing the distinct requirements imposed by mammalian hosts and bacterial targets. It also contributes to our rapidly expanding knowledge of inhibitors targeting gut microbial enzymes, an exciting frontier for chemical tool and therapeutic development.

## Acknowledgements

This work was funded by the Blavatnik Biomedical Accelerator at Harvard University and the Biocodex Microbiota Foundation. C.L. acknowledges support from the Jane Coffin Childs Memorial Fund for Medical Research. We thank François Lebreton and the Michael Gilmore lab of Mass Eye and Ear for generously providing bacterial strains, including the *E. faecalis* MMH594 Δ*tyrDC* strain. We thank Caroline Tanner and Ethan Brown for generously providing fecal samples from Parkinson’s disease patients. We acknowledge Minwoo Bae for critical reading of this manuscript. We acknowledge Xueyang Dong and Michelle Wang for assistance with cloning and protein purification, respectively.

## Synopsis

A known inhibitor of gut bacterial tyrosine decarboxylase enzymes is modified to disfavor metabolism by a human hydroxylase. This addresses a key limitation in applying gut bacterial insights to improve Parkinson’s disease therapy.

## Table of Contents Graphic

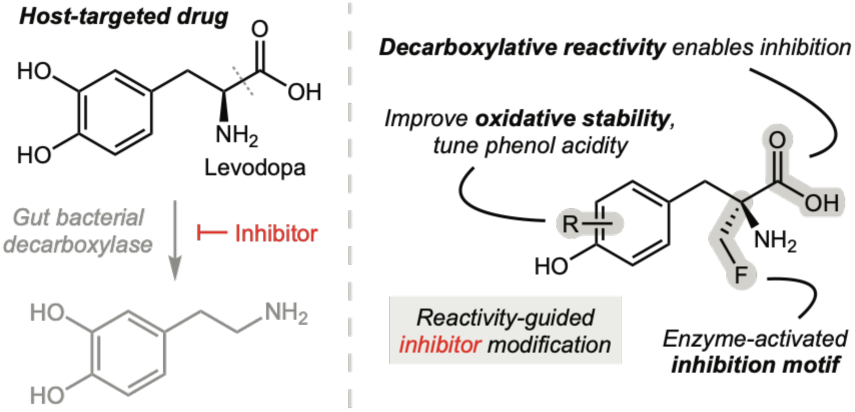

## Supporting Information

Experimental methods and materials, additional assay results, and synthetic details for reported compounds

