## Supplementary Material for "Inhibitors of gut bacterial L-dopa decarboxylation with reduced susceptibility to host metabolism"

### Table of Contents

Figure S1. Mechanism of L-dopa decarboxylation by PLP-dependent enzymes ........................ S3
Figure S2. Proposed mechanism for inhibition of TyrDC by AFMT ........................................ S3
Figure S3. Mechanism of tyrosine hydroxylation by TH ........................................................ S4
Figure S4. SDS-PAGE analysis of purified enzymes ............................................................... S4
Figure S5. Reduction of nitro-substituted amino acids by *E. faecalis* ....................................... S5
Figure S6. Inhibition of dopamine production in *E. faecalis* cultures ........................................ S5
Figure S7. Inhibition of dopamine production in assays with purified TyrDC ............................ S6
Figure S8. Inhibition of dopamine production in assays with purified AADC ........................... S7
Figure S9. Impact of inhibitors on dopamine production and growth of a panel of enterococci . S8
Figure S10. Inhibition of dopamine and *m*-tyramine production in fecal samples ...................... S9
Table S1. Suppliers and product numbers for aromatic amino acids ........................................ S10
Table S2. Suppliers and product numbers for general materials .............................................. S11
Table S3. UPLC–MS/MS parameters for compounds detected ................................................ S13
Table S4. Protein molecular weights and calculated extinction coefficients ............................ S14
General methods ..................................................................................................................... S14
UPLC–MS/MS methods .......................................................................................................... S14
Human TH expression and purification .................................................................................. S15
*E. faecalis* TyrDC expression and purification ........................................................................ S16
Human AADC expression and purification ............................................................................. S17
TH assays (Figures 1E, 3) ....................................................................................................... S17
Amino acid incubations with *E. faecalis* (Figures 2C, S5) ...................................................... S18
Inhibitor $EC_{50}$ assays with *E. faecalis* (Figures 4A, S6) .......................................................... S19
Inhibitor assays with purified TyrDC (Figures 4B, S7) ........................................................... S20
Inhibitor assays with purified AADC (Figure S8) ................................................................... S21
Inhibitor assays with a panel of enterococci (Figures 4C, S9) ................................................. S21
Inhibitor assays with human fecal samples (Figures 4D, S10) ................................................. S22
Synthetic methods ................................................................................................................... S24
Chromatograms and NMR spectra ........................................................................................... S39
References ............................................................................................................................... S44

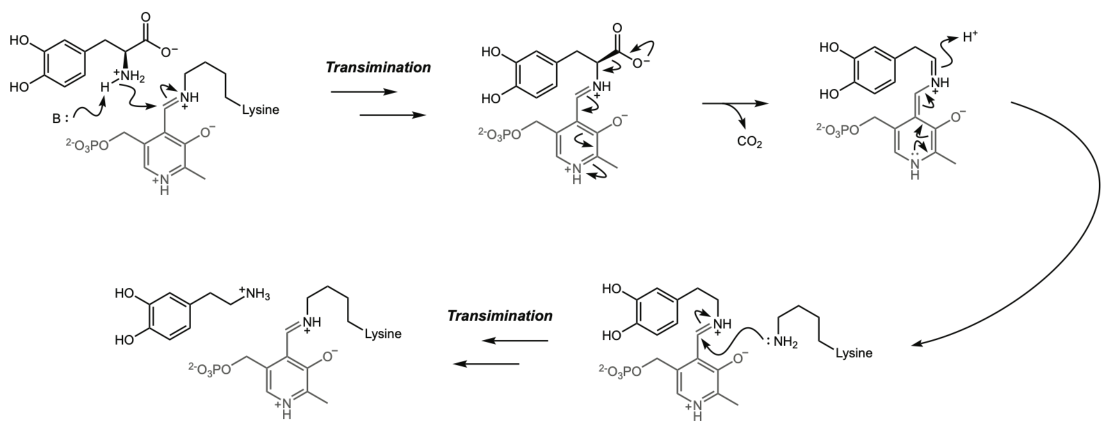


**Figure S1. Mechanism of L-dopa decarboxylation by PLP-dependent enzymes** The PLP cofactor rests as an internal aldiminium adduct with a lysine side chain. Catalysis initiates with transimination to connect the substrate and cofactor, followed by decarboxylation of the L-dopa substrate. The resulting quinonoid intermediate is protonated at the α-carbon, yielding an equivalent of dopamine bound to the PLP cofactor. A final transimination restores the internal aldiminium and releases the dopamine product.

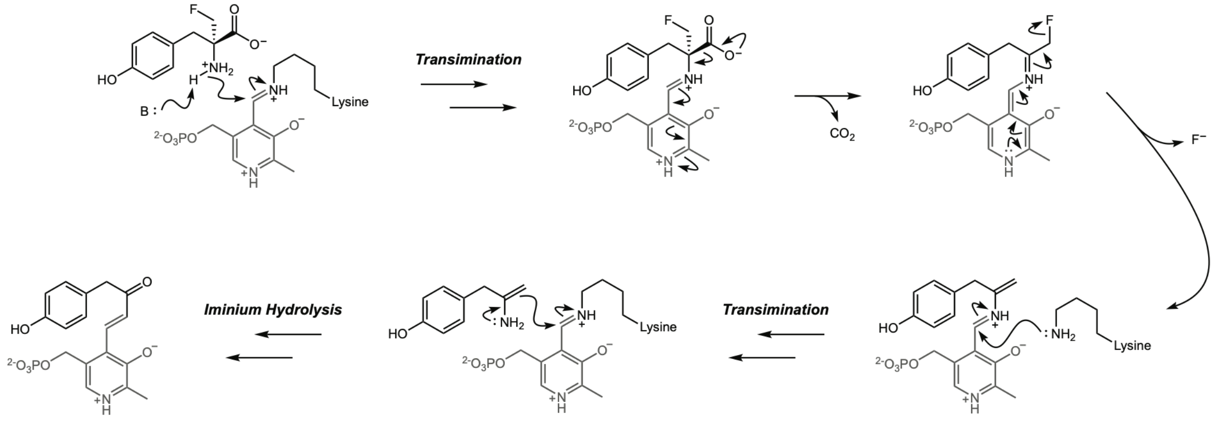


**Figure S2. Proposed mechanism for inhibition of TyrDC by AFMT** Catalysis initiates with engagement of the PLP cofactor by the inhibitor and subsequent decarboxylation. Instead of protonating at the α-carbon as happens with L-dopa, the quinonoid intermediate can lose an equivalent of fluoride, providing an enamine intermediate. The nucleophilic enamine can be released by transimination and react with the PLP cofactor, forming the adduct shown (detectable by mass spectrometry) and inactivating the enzyme.

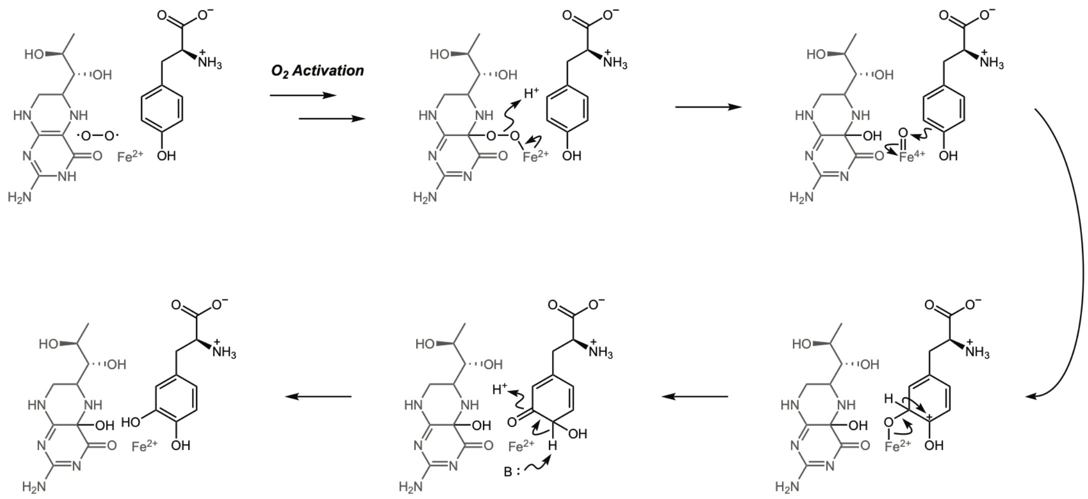


**Figure S3. Mechanism of tyrosine hydroxylation by TH** An equivalent of oxygen undergoes activation by the pterin co-substrate and Fe(II) center, allowing formation of a reactive Fe(IV) oxo species which is attacked by the aromatic ring of tyrosine. The cationic intermediate rearomatizes to provide the L-dopa product, which is released along with the oxidized pterin. For more information on this mechanism, see the provided references[1–4].

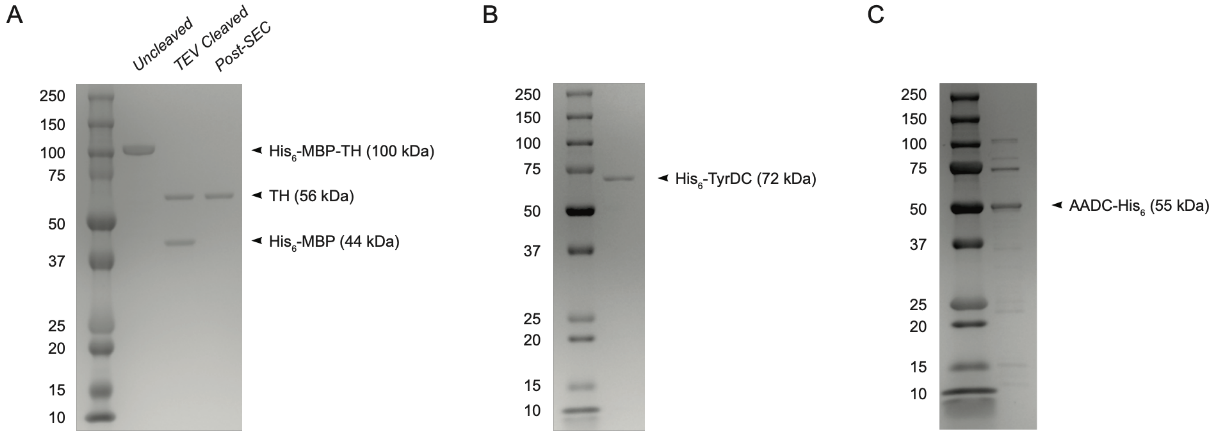


**Figure S4. SDS-PAGE analysis of purified enzymes** (*A*) Purified $His_6$-MBP-TH before and after cleavage by TEV protease, as well as the final purified TH after size exclusion chromatography. (*B*) Purified $His_6$-TyrDC. (*C*) Purified AADC-$His_6$. For all panels, the ladder bands are marked with nominal MW in kDa. See below for product and procedure details.

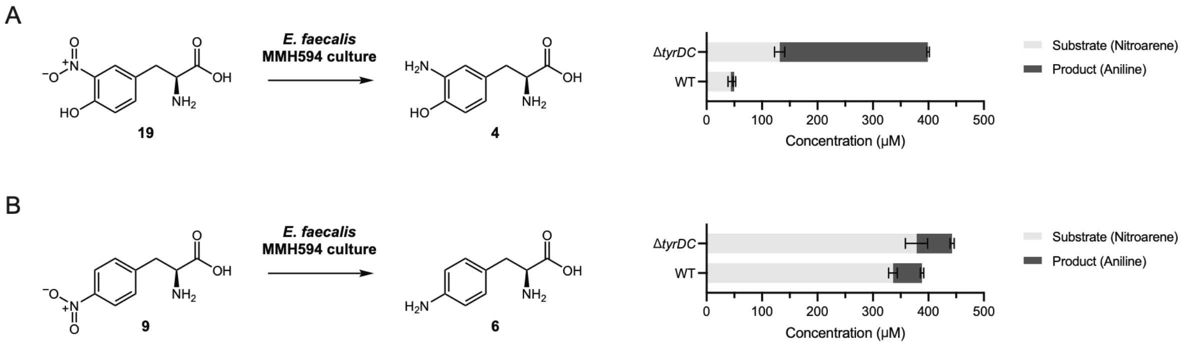


**Figure S5. Reduction of nitro-substituted amino acids by *E. faecalis*** (*A*) *E. faecalis* MMH594 WT or Δ*tyrDC* mutant strains were grown for 18 hours under anaerobic conditions with 500 µM **19** as a substrate. Post-incubation concentrations of the nitrobenzene substrate (**19**) and aniline reduction product (**4**) were determined using UPLC–MS/MS. (*B*) Same assay as in *A*, but using nitrobenzene substrate **9** to give aniline product **6**. For both panels, error bars show mean ± standard deviation of triplicate cultures grown in the same plate.

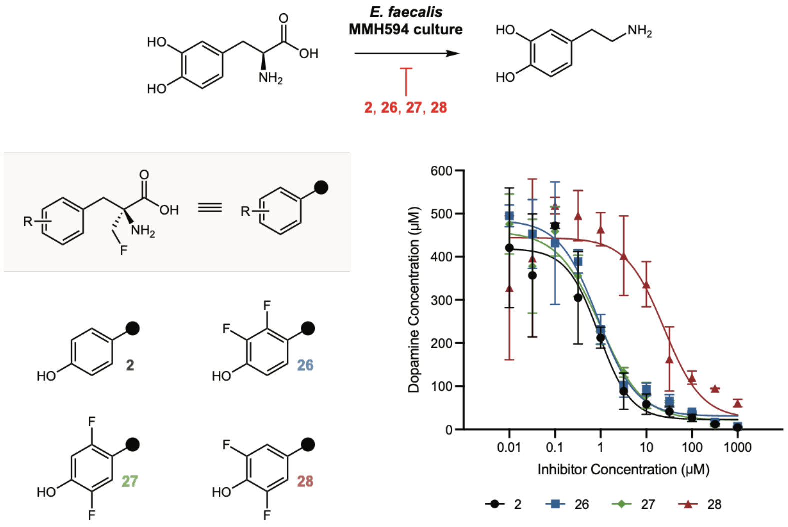


**Figure S6. Inhibition of dopamine production in *E. faecalis* cultures** *E. faecalis* MMH594 was grown anaerobically for 18 hours with 500 µM L-dopa and varying inhibitor concentrations. Dopamine production was quantified post-incubation using UPLC–MS/MS. Error bars show mean ± standard deviation of triplicate cultures grown in the same plate, and results are representative of two independent experiments conducted on different days.

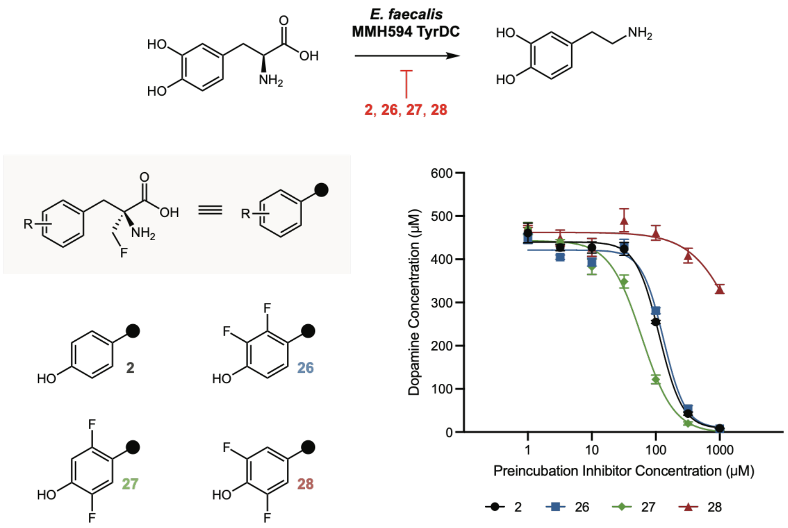


**Figure S7. Inhibition of dopamine production in assays with purified TyrDC** 1 µM purified TyrDC was preincubated for 20 minutes with 200 µM PLP and varying inhibitor concentrations at room temperature in pH 7 HEPES buffer. The preincubations were then diluted 10X in a solution of substrate to give assay concentrations of 500 µM L-dopa, 0.1 µM TyrDC, 20 µM PLP, and varying inhibitor concentrations in pH 5.5 sodium acetate buffer. Dopamine production after 20 minutes of reaction was quantified using UPLC–MS/MS. Note that inhibitor concentrations used for preincubation are plotted. Error bars show mean ± standard deviation of triplicate assays run in the same plate, and results are representative of two independent experiments conducted on different days.

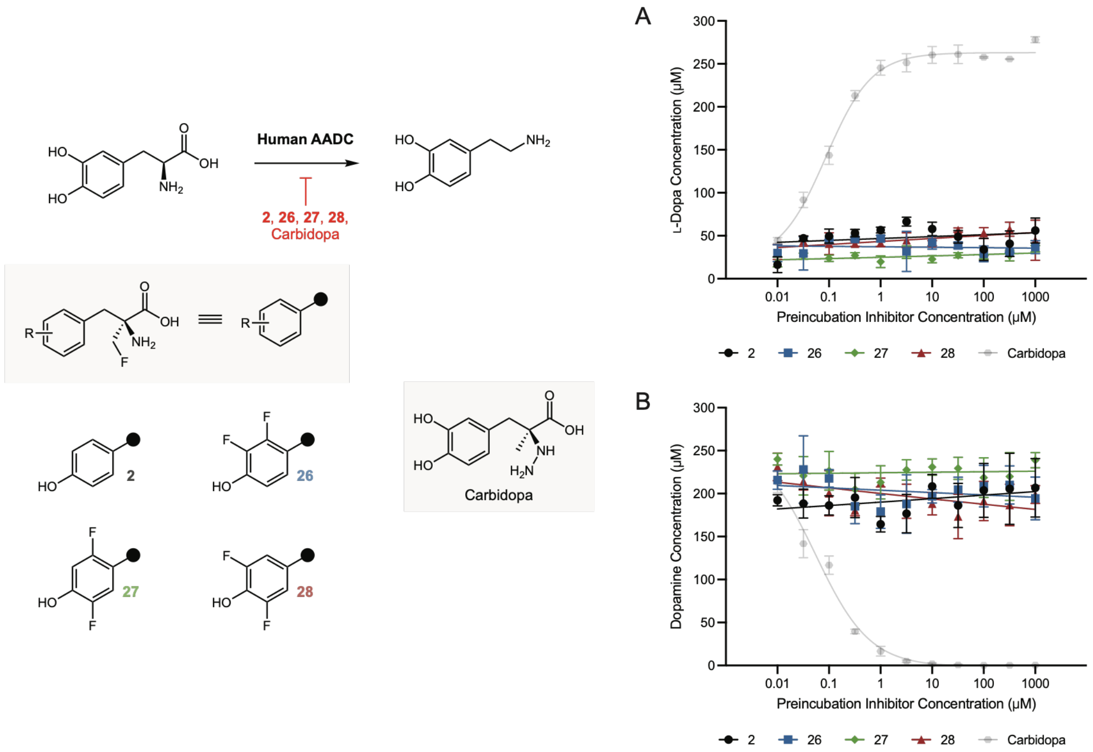


**Figure S8. Inhibition of dopamine production in assays with purified AADC** (*A*) 2 µM purified AADC was preincubated for 20 minutes with 25 µM PLP and varying inhibitor concentrations at room temperature in pH 7 HEPES buffer. The preincubations were then diluted 10X in a solution of substrate to give assay concentrations of 250 µM L-dopa, 0.2 µM AADC, 2.5 µM PLP, and varying inhibitor concentrations in pH 7.4 potassium phosphate buffer. Remaining substrate after 30 minutes of reaction at 37 °C was quantified using UPLC–MS/MS. (*B*) Dopamine production in the same assay. Note that inhibitor concentrations used for preincubation are plotted. Error bars show mean ± standard deviation of triplicate assays run in the same plate, and results are representative of two independent experiments conducted on different days.

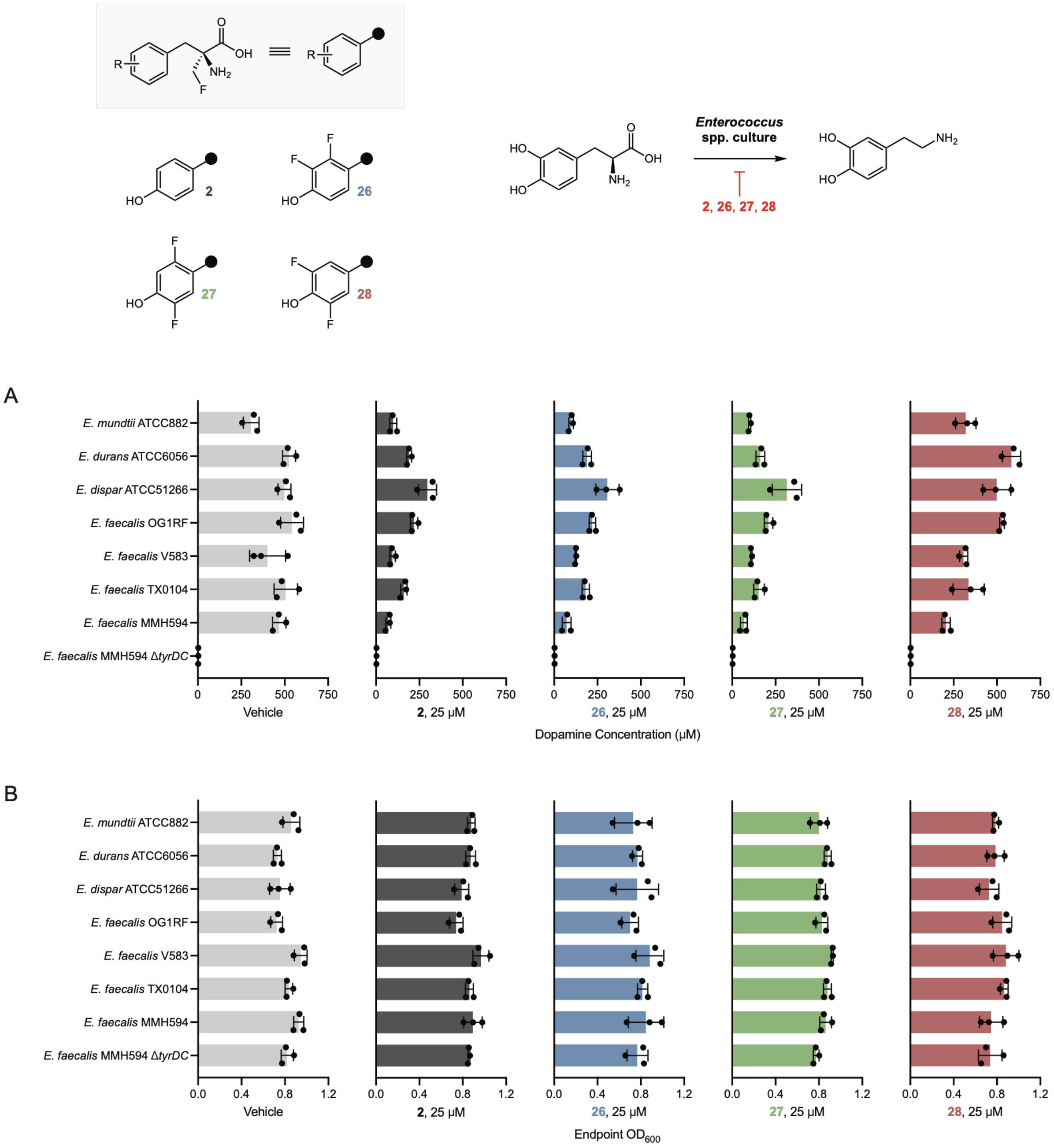


**Figure S9. Impact of inhibitors on dopamine production and growth of a panel of enterococci** (*A*) Each strain was grown anaerobically for 18 hours in the presence of 500 µM L-dopa and 25 µM inhibitor, after which dopamine production was quantified using UPLC–MS/MS. (*B*) Post-incubation $OD_{600}$ values for all strain/inhibitor combinations. For both panels, error bars show mean ± standard deviation of triplicate cultures run in the same plate, and results are representative of two independent experiments conducted on different days.

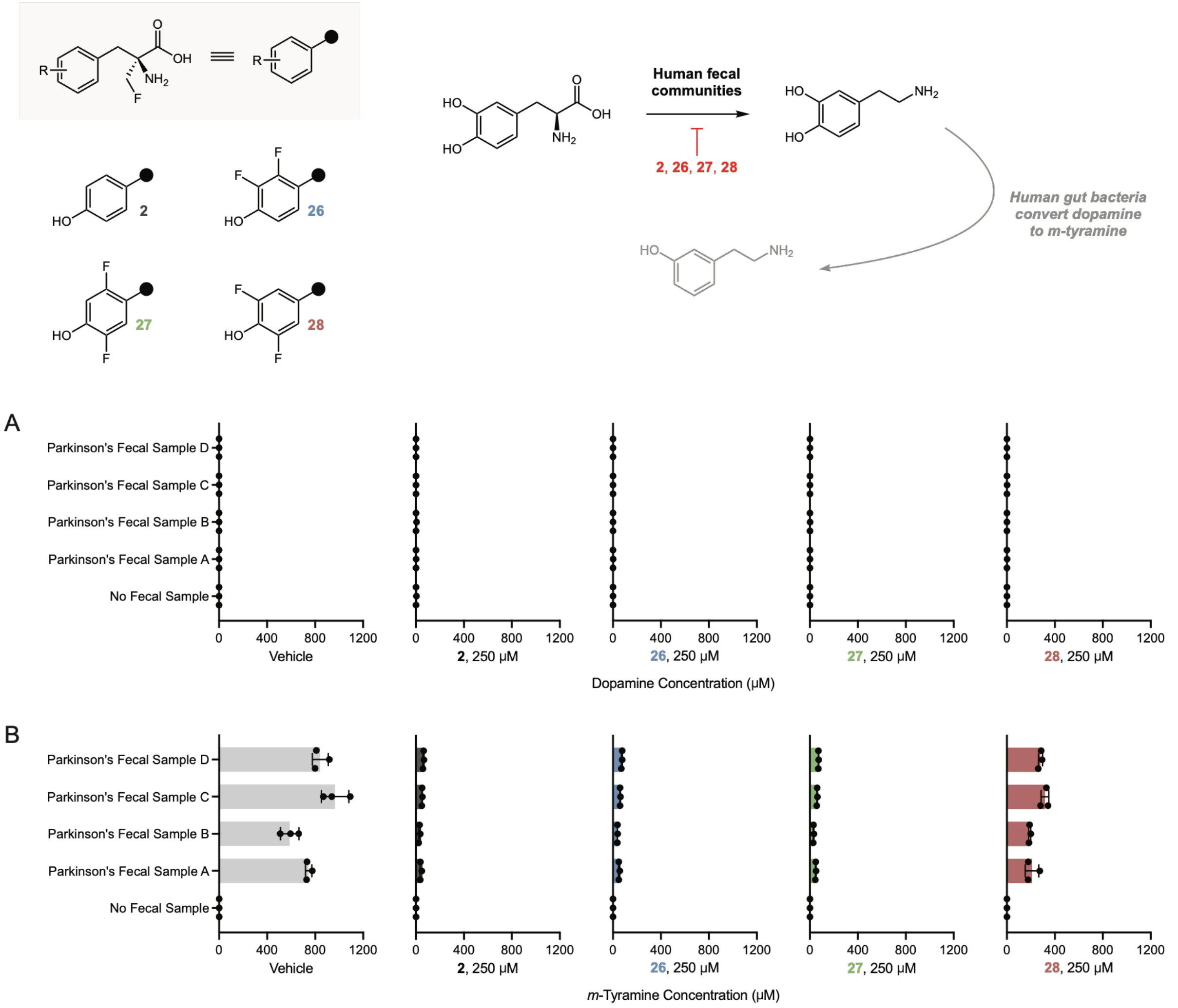


**Figure S10. Inhibition of dopamine and *m*-tyramine production in fecal samples** (*A*) Fecal communities were grown for 40 hours in the presence of 1000 µM L-dopa and 250 µM inhibitor, after which dopamine production was quantified using UPLC–MS/MS. (*B*) *m*-Tyramine production in the same assay. All four samples were provided by distinct donors. For both panels, error bars show mean ± standard deviation of triplicate community cultures grown in the same plate.

**Table S1. Suppliers and product numbers for aromatic amino acids**

| Compound Number | Compound Name | Supplier | Product Number |
|---|---|---|---|
| | L-Tyrosine hydrochloride | Chem-Impex | 00303 |
| **1** | L-Dopa | Sigma-Aldrich | D9628 |
| **4** | 3-Amino-L-tyrosine dihydrochloride | Oakwood Chemical | M03158 |
| **5** | L-m-tyrosine | Alfa Aesar | H27224 |
| **6** | 4-Amino-L-phenylalanine | Chem-Impex | 02594 |
| **7** | (*S*)-2-Amino-3-(4-(hydroxymethyl)phenyl)propanoic acid hydrochloride | AstaTech | AT22619 |
| **8** | 4-Carboxy-L-phenylalanine | Chem-Impex | 07030 |
| **9** | 4-Nitro-L-phenylalanine | Oakwood Chemical | 036596 |
| **10** | 3-Fluoro-L-phenylalanine | Chem-Impex | 02571 |
| **11** | 4-Fluoro-L-phenylalanine hydrochloride | Oakwood Chemical | 008572 |
| **12** | L-3,4-Difluorophenylalanine | Chem-Impex | 04117 |
| **13** | (*S*)-2-Amino-3-(2-fluoro-4-hydroxyphenyl)propanoic acid hydrochloride | AstaTech | Y13603 |
| **14** | (*S*)-2-Amino-3-(2-chloro-4-hydroxyphenyl)propanoic acid hydrochloride | AstaTech | Y14207 |
| **15** | 3-Fluoro-L-tyrosine | AstaTech | 61380 |
| **16** | 3-Chlorotyrosine | Oakwood Chemical | M03243 |
| **17** | 3-Bromo-L-tyrosine | AstaTech | A11383 |
| **18** | H-Tyr(3-I)-OH | Ambeed | A190487 |
| **19** | 3-Nitro-L-tyrosine | Ambeed | A708002 |
| **20** | 3,5-Difluoro-L-tyrosine | AstaTech | 98380 |
| **21** | 3-(2’-Pyridyl)-L-alanine | Chem-Impex | 06211 |
| **22** | 3-(3’-Pyridyl)-L-alanine | Chem-Impex | 02853 |
| **23** | 3-(4’-Pyridyl)-L-alanine | Chem-Impex | 06924 |
| **24** | (2*S*)-2-Amino-3-(6-hydroxypyridin-3-yl)propanoic acid | ChemSpace | CSC015411575 |
| **25** | 5-Hydroxy-L-tryptophan | Sigma-Aldrich | H9772 |

**Table S2. Suppliers and product numbers for general materials**

| Product Name | Supplier | Product Number |
|---|---|---|
| Glycerol | Sigma-Aldrich | G5516 |
| Kanamycin Sulfate, Ultra Pure Grade | VWR | 0408-EU |
| IPTG, Ultra Pure Grade | Teknova, VWR | I3325, 0487 |
| LB Broth Powder (Lennox Broth) | RPI | L24060 |
| Difco LB Agar, Lennox | BD | 240110 |
| BBL Brain Heart Infusion | BD | 211059 |
| Anaerobic Hungate Culture Tubes | Chemglass | CLS-4208 |
| BD GasPak EZ anaerobe pouch system | BD | 260683 |
| 96-well Clear Flat Bottom Polystyrene TC-treated Microplate | Corning | 3595 |
| qPCR Tube Strips with Individually-Attached Caps | VWR | 93001-0118 |
| Microseal 'F' PCR Plate Seal | Bio-Rad | MSF1001 |
| Syringe Filter | VWR | 76479-024 |
| Q5 High-Fidelity 2X Master Mix | NEB | M0492S |
| NdeI | NEB | R0111S |
| XhoI | NEB | R0146S |
| rCutSmart Buffer | NEB | B6004S |
| Ni-NTA Agarose | Qiagen | 30250 |
| Amicon Ultra Centrifugal Filter, 50 kDa MWCO | Millipore | UFC9050 |
| Amicon Ultra Centrifugal Filter, 30 kDa MWCO | Millipore | UFC9030 |
| Amicon Ultra Centrifugal Filter, 10 kDa MWCO | Millipore | UFC9010 |
| PD-10 desalting columns packed with Sephadex G-25 resin | Cytiva | 17085101 |
| TEV Protease | NEB | P8112S |

| | | |
|---|---|---|
| Novex Tris-Glycine Mini Protein Gels, 10-20%, 1.0 mm, WedgeWell format | ThermoFisher | XP10205BOX |
| 10x Tris/Glycine/SDS | Bio-Rad | 1610772 |
| 2x Laemmli Sample Buffer | Bio-Rad | 1610737 |
| 2-Mercaptoethanol | Sigma-Aldrich | M3148 |
| Precision Plus Protein Dual Color Standards | Bio-Rad | 1610374 |
| Precision Plus Protein Dual Xtra Prestained Protein Standards | Bio-Rad | 1610377 |
| InstantBlue Coomassie Protein Stain | Abcam | ab119211 |
| Sodium chloride | Sigma-Aldrich | S9888 |
| HEPES | Sigma-Aldrich | RDD002 |
| Sodium acetate | Sigma-Aldrich | S2889 |
| Potassium Phosphate, Monobasic | RPI | P41200 |
| Potassium phosphate dibasic | Sigma-Aldrich | 795496 |
| Dopamine hydrochloride | Supelco | PHR1090 |
| Pyridoxal 5'-phosphate monohydrate | Sigma-Aldrich | 82870 |
| (6*R*)-5,6,7,8-Tetrahydrobiopterin dihydrochloride | Sigma-Aldrich | T4425 |
| Catalase from bovine liver | Sigma-Aldrich | C9322 |
| Ammonium iron(II) sulfate hexahydrate | Sigma-Aldrich | F1543 |
| L-Dopa-(*phenyl*-$d_3$) | Sigma-Aldrich | 333786 |
| 1,4-Dithiothreitol | Roche | 10708984001 |
| Formic acid 98% - 100% | Supelco | 5.33002 |
| Acetonitrile | Sigma-Aldrich | 900667 |

**Table S3. UPLC–MS/MS parameters for compounds detected**

| Compound Name or Number | Retention Time (min) | Parent Ion (m/z) | Daughter Ion (m/z) | Cone Voltage (V) | Collision Voltage (V) |
|---|---|---|---|---|---|
| L-Tyrosine | 0.90 | 182.1058 | 136.1089 | 16 | 14 |
| Dopamine | 0.70 | 154.1109 | 91.0764 | 16 | 18 |
| *meta*-Tyramine | 1.40 | 138.1119 | 121.1143 | 28 | 10 |
| L-Dopa-(*phenyl*-$d_3$) | 0.60 | 201.1195 | 154.4441 | 4 | 14 |
| **1** | 0.60 | 198.1645 | 152.0929 | 32 | 14 |
| **2** | 1.20 | 213.9843 | 168.1005 | 20 | 12 |
| **3** | 0.80 | 229.9843 | 184.0941 | 24 | 14 |
| **4** | 0.50 | 197.2396 | 180.0161 | 30 | 8 |
| **5** | 1.10 | 182.2296 | 136.0560 | 28 | 12 |
| **6** | 0.50 | 181.2496 | 164.0205 | 26 | 8 |
| **7** | 1.35 | 196.2496 | 150.0251 | 32 | 12 |
| **8** | 1.80 | 210.2296 | 164.0722 | 30 | 12 |
| **9** | 2.30 | 211.2196 | 119.0117 | 34 | 24 |
| **10** | 2.25 | 184.2296 | 118.0322 | 30 | 20 |
| **11** | 2.25 | 184.2296 | 118.0322 | 30 | 22 |
| **12** | 2.30 | 202.2196 | 136.0074 | 34 | 22 |
| **13** | 1.30 | 200.0919 | 154.0919 | 32 | 14 |
| **14** | 2.25 | 216.0619 | 170.1094 | 38 | 14 |
| **15** | 1.15 | 200.0919 | 154.0936 | 40 | 14 |
| **16** | 2.25 | 216.0619 | 170.1076 | 32 | 12 |
| **17** | 2.30 | 260.0119 | 214.0455 | 36 | 14 |
| **18** | 2.35 | 308.0019 | 262.0342 | 22 | 14 |
| **19** | 2.30 | 227.0919 | 181.0791 | 36 | 12 |
| **20** | 1.45 | 218.0919 | 172.0881 | 36 | 12 |
| **21** | 0.50 | 167.2296 | 121.0433 | 40 | 14 |
| **22** | 0.50 | 167.2296 | 106.0160 | 32 | 14 |
| **23** | 0.45 | 167.2296 | 149.9933 | 30 | 12 |
| **24** | 0.50 | 183.2296 | 109.0730 | 40 | 20 |
| **25** | 2.00 | 220.9843 | 204.0927 | 30 | 10 |
| **26** | 2.20 | 250.0281 | 204.0750 | 34 | 14 |
| **27** | 2.10 | 250.0281 | 204.0750 | 2 | 14 |
| **28** | 2.10 | 250.0281 | 204.0750 | 6 | 14 |

**Table S4. Protein molecular weights and calculated extinction coefficients**

| Construct | MW (kDa) | Extinction Coefficient |
|---|---|---|
| $His_6$-MBP-TH | 99.755 | 110,045 |
| $His_6$-MBP *[TEV cleaved]* | 44.104 | 69,330 |
| TH *[TEV cleaved]* | 55.669 | 40,715 |
| $His_6$-TyrDC | 72.393 | 114,140 |
| AADC-$His_6$ | 55.446 | 69,620 |

**General methods**

TH buffer consists of 50 mM HEPES and 100 mM NaCl at pH 7. TyrDC buffer consists of 50 mM Tris and 250 mM NaCl at pH 8. AADC buffer consists of 20 mM sodium phosphate and 500 mM NaCl at pH 7.4. KPhos buffer consists of 100 mM potassium phosphate at pH 7.4. NaOAc buffer consists of 200 mM sodium acetate at pH 5.5. Centrifugation steps were performed with an Eppendorf 5810R or Sorvall Lynx 6000 centrifuge. Protein concentrations were determined using a Nanodrop 2000 spectrophotometer and extinction coefficients calculated with Geneious Prime. Protein loading in SDS-PAGE gels was 2 µg per lane, and electrophoresis was conducted using a Bio-Rad PowerPac Basic power supply. Gels were run at 100 V for 10 minutes, then 150 V for 45 minutes, followed by staining with InstantBlue and destaining as necessary for clear imaging. Precision Plus Protein Dual Color or Dual Xtra standards were included on each protein gel to allow MW comparison.

Bacterial strains were obtained commercially or were generously provided by François Lebreton and the Michael Gilmore lab of Mass Eye and Ear. Plate-based $OD_{600}$ measurements were made with a BioTek Synergy Neo2 plate reader. All anaerobic bacterial growth experiments were conducted in a Coy Laboratory Products anaerobic chamber under an atmosphere of 5% hydrogen, 10% carbon dioxide, and balance nitrogen. Plastics and media used in anaerobic experiments were brought into the chamber the day before use to allow deoxygenation. We note that solubility is a general challenge for the compounds of interest, and many required sustained sonication and/or vortexing to dissolve. L-Dopa solutions were made on the day of assays to avoid oxidation.

**UPLC–MS/MS methods**

All UPLC–MS/MS analysis was performed using a Waters Acquity UPLC H-Class PLUS System and a Waters Xevo TQ-S mass spectrometer. A CORTECS T3 2.7 µm (2.1 x 100 mm) column was used with mobile phase A 0.1% formic acid in water and mobile phase B 0.1% formic acid in acetonitrile. The gradient used was as follows: 0-1 min 0% B isocratic, 1-2 min 0-90% B, 2-2.5 min 90% B isocratic, 2.5-2.75 min 90-0% B, 2.75-3.4 min 0% B isocratic. Column temperature was 40 °C, flow rate was 0.5 mL/min, and injection volume was 2 µL. All compounds were detected in positive mode using electrospray ionization and multiple reaction monitoring. Mass transitions, cone voltages, collision voltages, and retention times for each compound are reported in Supplementary Table 3. Standard curves of each compound were generated on the day of analysis to allow quantitation. Data was processed using TargetLynx software from Waters.

**Human TH expression and purification**

Expression and purification followed a protocol adapted from Higgins et al.[5] and Bueno-Carrasco et al.[6]

cDNA encoding human TH was purchased from Sino Biological in a cloning vector (product HG10684-M). The TH insert was amplified from the cloning vector using PCR, as was a maltose-binding protein (MBP) insert from an in-house plasmid that included a linker and TEV cleavage site. An empty pET-28a vector was digested using NdeI and XhoI, then combined with the two inserts by Gibson assembly, yielding a $His_6$-MBP-TH fusion construct with a TEV cleavage site between MBP and TH. After confirmation by Sanger sequencing, the pET-28a vector encoding $His_6$-MBP-TH was transformed into *E. coli* BL21(DE3).

A starter culture was grown from the glycerol stock of the transformed strain in 6 mL of LB medium with 50 µg/mL kanamycin sulfate for 18 hours at 37 °C with shaking at 200 rpm in an Innova 42R incubator shaker. The starter culture was used to inoculate 4 L of LB medium (split into two flasks) with 50 µg/mL kanamycin sulfate. The expression cultures were grown at 37 °C with shaking at 180 rpm in an Innova S44i incubator shaker until the $OD_{600}$ reached 0.5, at which point isopropyl β-D-1-thiogalactopyranoside (IPTG, final concentration 200 µM) and ferrous ammonium sulfate hexahydrate (final concentration 100 µM) were added as sterile solutions in ultrapure water. The expression cultures were then kept at 25 °C with shaking at 180 rpm in an Innova S44i incubator shaker for 17 hours.

Cultures were harvested by centrifugation for 12 minutes at 6700 rcf and 4 °C. The combined pellet was resuspended in 30 mL of lysis buffer and lysed by sonication using a Branson 450 Digital Sonifier (3 minute sonication time, 2 seconds on 10 seconds off at 25% amplitude with a 0.5 inch horn, performed 3 times with sample mixing in between). Lysis buffer was made by dissolving one EDTA-free Pierce protease inhibitor tablet in 50 mL of TH buffer. Sonication was performed at 4 °C, as were all remaining steps in protein purification. The lysate was clarified by centrifugation at 16000 rcf for 20 minutes.

The clarified lysate was incubated for 30 minutes with 6 mL of Ni-NTA resin before loading onto a column for elution. The resin was eluted with 10 mL aliquots of TH buffer containing the following imidazole concentrations (in order): 2 x 75 mM, 2 x 100 mM, 125 mM, 150 mM, 200 mM, 2 x 250 mM. All fractions of 100 mM or higher imidazole concentration were combined and concentrated to a final volume of 1.6 mL using spin concentrators with a 50 kDa MW cutoff. The concentrated protein was desalted using a PD-10 desalting column following manufacturer instructions.

The $His_6$-MBP tag was removed by TEV digestion as follows. 1 mL of 10X reaction buffer, 100 µL of TEV protease, and 474 µL of $His_6$-MBP-TH were diluted with ultrapure water to a final volume of 10 mL (four separate digests were set up following this ratio). Digests were left for 38 hours before combining and concentrating to a final volume of 2 mL using spin concentrators with a 30 kDa MW cutoff. The mixture was separated using a Cytiva ÄKTApure M 25 FPLC with a HiLoad 16/600 Superdex 200 pg size exclusion column. 1.3 column volumes of TH buffer were used for isocratic elution, with fractions collected every 4 mL after 0.25 column volumes.

Fractions were analyzed by SDS-PAGE, and those containing purified TH were combined and concentrated to a final volume of 5 mL using spin concentrators with a 30 kDa MW cutoff. 200 µL aliquots were flash-frozen in liquid nitrogen and stored at –80 °C until use. Calculated yield of purified TH was 5.77 mg from 4 L of expression culture.

***E. faecalis* TyrDC expression and purification**

Expression and purification followed a similar protocol to that reported previously, using the same pET-28a $His_6$-TyrDC construct and *E. coli* BL21(DE3) expression strain[7].

A starter culture was grown from the glycerol stock of the expression strain in 6 mL of LB medium with 50 µg/mL kanamycin sulfate for 18 hours at 37 °C with shaking at 200 rpm in an Innova 42R incubator shaker. 3 mL of starter culture was used to inoculate 2 L of LB medium with 50 µg/mL kanamycin sulfate. The expression culture was grown at 37 °C with shaking at 180 rpm in an Innova 42R incubator shaker until the $OD_{600}$ reached 0.35, at which point IPTG (final concentration 200 µM) was added as a sterile solution in ultrapure water. The expression culture was then kept at 18 °C with shaking at 180 rpm in an Innova 42R incubator shaker for 18 hours.

The culture was harvested by centrifugation for 12 minutes at 6700 rcf and 4 °C. The pellet was stored at –80 °C for less than 4 weeks before protein purification. After thawing, the pellet was resuspended in 30 mL of lysis buffer and lysed by sonication using a Branson 450 Digital Sonifier (3 minute sonication time, 2 seconds on 10 seconds off at 25% amplitude with a 0.5 inch horn, performed 2 times with sample mixing in between). Lysis buffer was made by dissolving one Pierce protease inhibitor tablet in 50 mL of TyrDC buffer. Sonication was performed at 4 °C, as were all remaining steps in protein purification. The lysate was clarified by centrifugation at 16000 rcf for 20 minutes.

The clarified lysate was incubated for 2 hours with 3 mL of Ni-NTA resin before loading onto a column for elution. The resin was eluted with 10 mL aliquots of TyrDC buffer containing the following imidazole concentrations (in order): 2 x 75 mM, 2 x 100 mM, 125 mM, 150 mM, 200 mM, 2 x 250 mM. All fractions of 100 mM or higher imidazole concentration were combined and concentrated to a final volume of 2 mL using spin concentrators with a 30 kDa MW cutoff. The concentrated protein was desalted using a PD-10 desalting column following manufacturer instructions.

250 µL aliquots were flash-frozen in liquid nitrogen and stored at –80 °C until use. Calculated yield of purified TyrDC was 17.11 mg from 2 L of expression culture.

**Human AADC expression and purification**

Expression and purification followed a protocol adapted from our prior paper[7] and literature examples, including Montioli et al.[8] and van Kessel et al.[9] A pET-28a AADC-$His_6$ construct was purchased from Twist Bioscience's clonal genes service and transformed into *E. coli* BL21(DE3).

A starter culture was grown from the glycerol stock of the expression strain in 6 mL of LB medium with 50 µg/mL kanamycin sulfate for 18 hours at 37 °C with shaking at 200 rpm in an Innova 42R incubator shaker. The starter culture was used to inoculate 4 L of LB medium (split into two flasks) with 50 µg/mL kanamycin sulfate. The expression cultures were grown at 37 °C with shaking at 180 rpm in an Innova 42R incubator shaker until the $OD_{600}$ reached 0.5, at which point IPTG (final concentration 200 µM) was added as a sterile solution in ultrapure water. The expression cultures were then kept at 18 °C with shaking at 180 rpm in an Innova 42R incubator shaker for 18 hours.

Cultures were harvested by centrifugation for 12 minutes at 6700 rcf and 4 °C. The combined pellet was resuspended in 25 mL of lysis buffer and lysed by sonication using a Branson 450 Digital Sonifier (3 minute sonication time, 2 seconds on 10 seconds off at 25% amplitude with a 0.5 inch horn, performed 3 times with sample mixing in between). Lysis buffer was made by dissolving one Pierce protease inhibitor tablet in 50 mL of AADC buffer with 20 mM imidazole and 50 µM PLP. Sonication was performed at 4 °C, as were all remaining steps in protein purification. The lysate was clarified by centrifugation at 16000 rcf for 20 minutes.

The clarified lysate was incubated for one hour with 5.5 mL of Ni-NTA resin before loading onto a column for elution. The resin was eluted with 10 mL aliquots of AADC buffer containing the following imidazole concentrations (in order): 2 x 75 mM, 2 x 100 mM, 125 mM, 150 mM, 200 mM, 2 x 250 mM. The last six fractions were combined and concentrated using spin concentrators with a 10 kDa MW cutoff. Desalting was accomplished through three rounds of concentration and dilution with KPhos buffer containing 100 µM PLP and 10% glycerol by volume.

400 µL aliquots were flash-frozen in liquid nitrogen and stored at –80 °C until use. Calculated yield of purified AADC was 4.76 mg from 4 L of expression culture.

**TH assays (Figures 1E, 3)**

TH activity assays followed a protocol adapted from Bueno-Carrasco et al.[6]

The concentrations of each assay component were as follows: 0.5 µM TH, 50 µM substrate, 0.1 mg/mL bovine liver catalase, 10 µM ferrous ammonium sulfate hexahydrate, 300 µM tetrahydrobiopterin, and 5 mM 1,4-dithiothreitol (DTT). TH buffer was used for all solutions and dilutions (prior to quenching in organic solvent). Solutions of catalase, DTT, and ferrous ammonium sulfate hexahydrate were made fresh before assay setup. Tetrahydrobiopterin was solubilized immediately before addition. Control assays without TH were prepared with 750 µL of buffer used to substitute for 750 µL of TH solution. The protocol is detailed below.

A flash-frozen TH aliquot was thawed at 4 °C before warming to room temperature. The aliquot was then diluted with buffer to yield a 2 μM TH solution. 750 μL of the 2 μM TH solution was combined with 300 μL of bovine liver catalase (1 mg/mL), 30 μL of ferrous ammonium sulfate hexahydrate (1 mM), and 1170 μL of buffer before mixing well with a micropipette. 225 μL aliquots of this mastermix were combined with 15 μL each of the appropriate substrate solution (1 mM, either amino acids or AFMT analogs) and mixed before distributing across triplicate PCR tubes in 80 μL aliquots. A solution containing and DTT (25 mM) and tetrahydrobiopterin (1.5 mM) was prepared after distribution of the enzyme/substrate mix into PCR tubes. The PCR tubes were incubated for 1 minute at 37 °C before the reaction was initiated by adding 20 μL of the tetrahydrobiopterin/DTT mix. The reaction mixtures were mixed with a multichannel pipette set to 50 μL before incubation at 37 °C. Reaction mixtures were mixed again and aliquots were quenched by 10X dilution (20 μL in 180 μL) in mass spectrometry grade methanol or acetonitrile at the appropriate timepoints, followed by brief storage at –80 °C.

For assays with tyrosine analogs (Figure 3A), aliquots were quenched in methanol after 1 hour. For the assay with AFMT shown in Figure 1E, aliquots were quenched in methanol after 2 hours. In both cases, the quenched aliquots were kept at –80 °C for less than 2 hours, after which they were warmed to room temperature and mixed with a multichannel pipette set to 200 μL before centrifugation for 30 minutes at 3220 rcf. The centrifuged samples were then diluted 5X in ultrapure water (40 μL in 160 μL) and mixed with a multichannel pipette before UPLC–MS/MS analysis.

For assays with AFMT derivatives (Figure 3B), aliquots were quenched in acetonitrile after 2 hours. The quenched aliquots were kept at –80 °C for less than 2 hours, after which they were warmed to room temperature and mixed with a multichannel pipette set to 200 μL before centrifugation for 30 minutes at 3220 rcf. The centrifuged samples were then diluted 10X in ultrapure water containing 5.55 μM L-dopa-(*phenyl*-$d_3$) as an internal standard (20 μL in 180 μL) and mixed with a multichannel pipette set to 150 μL before UPLC–MS/MS analysis.

For concentration normalization to no-enzyme controls, concentration in each TH-containing reaction mixture was divided by the mean of concentrations for all three no-enzyme reaction mixtures at the corresponding timepoint. The normalized data points were then used for plotting.

**Amino acid incubations with *E. faecalis* (Figures 2C, S5)**

The concentration of amino acid substrates in assays was 500 μM. Between 6 and 10 substrates were analyzed per experiment. Consumables needed for assay setup were brought into an anaerobic chamber one day before use.

Starter cultures of *E. faecalis* MMH594 WT and Δ*tyrDC* mutant strains were inoculated from glycerol stocks in the anaerobic chamber using 6 mL of BHI medium in Hungate tubes. The tubes were sealed, brought out of the chamber, and cultured for 18 hours at 37 °C without shaking.

The next day, 555 μM solutions of each substrate were made in the chamber by dissolving an appropriate amount of solid in 45 mL of BHI medium (this step requires sustained vortexing for some substrates). The pH of the medium was not controlled for perturbations introduced by substrate dissolution. These substrate solutions were distributed into separate wells of a 96-well plate in 180 μL aliquots. The *E. faecalis* starter cultures were diluted 10X into the substrate solutions (20 μL into 180 μL) using a multichannel pipette, after which the plate was sealed using a foil plate seal and moved into a GasPak anaerobic pouch. Triplicate wells were included for each substrate/strain combination. The anaerobic pouch was removed from the chamber and placed in a 37 °C incubator for 18 hours without shaking.

After incubation, the plate was frozen at –80 °C for less than 12 hours before warming to room temperature. The samples were mixed with a multichannel pipette set to 100 μL before 25X dilution (8 μL in 192 μL) in mass spectrometry grade methanol. The methanol dilutions were mixed with a multichannel pipette set to 100 μL then centrifuged for 5 minutes at 3220 rcf. The centrifuged samples were then diluted 20X or 40X in ultrapure water (10 or 5 μL in 190 or 195 μL, respectively) and mixed with a multichannel pipette before UPLC–MS/MS analysis.

**Inhibitor $EC_{50}$ assays with *E. faecalis* (Figures 4A, S6)**

The concentration of L-dopa in assays was 500 μM and the concentration of inhibitors in assays varied from 0.01-1000 μM. Inhibitors were dissolved to a concentration of 1.5 mM in BHI medium with vigorous shaking and sonication, after which the solutions were sterilized using 0.22 μm filters. The sterilized inhibitor stocks were diluted (using sterile technique) with more BHI medium to provide the desired inhibitor concentrations in 450 μL aliquots. These aliquots were brought into an anaerobic chamber one day before use, as were the consumables needed for assay setup. A starter culture of *E. faecalis* MMH594 was inoculated from a glycerol stock in the anaerobic chamber using 8 mL of BHI medium in a Hungate tube. The tube was sealed, brought out of the chamber, and cultured for 18 hours at 37 °C without shaking.

The next day, a 2.22 mM L-dopa solution was made in the chamber by dissolving 13.1 mg of L-dopa in 30 mL of BHI medium (this step requires sustained vortexing). 150 μL of the 2.22 mM L-dopa solution was added to each 450 μL inhibitor aliquot that had equilibrated overnight, providing solutions containing 555 μM L-dopa and varied inhibitor concentrations. These solutions were mixed by vortexing, then distributed into 96-well plates in 180 μL aliquots. The *E. faecalis* starter culture was diluted 10X into the inhibitor/L-dopa solutions (20 μL into 180 μL) using a multichannel pipette, after which the plates were sealed using foil plate seals and moved into a GasPak anaerobic pouch. Triplicate wells were included for each inhibitor concentration, as well as for control assays containing the WT bacterium or a Δ*tyrDC* mutant strain with no inhibitor. The anaerobic pouch was removed from the chamber and placed in a 37 °C incubator for 18 hours without shaking.

After incubation, the plates were frozen at –80 °C until UPLC–MS/MS analysis (within 5 days). When ready, the plates were warmed to room temperature and mixed with a multichannel pipette set to 100 μL before 10X dilution (20 μL in 180 μL) in mass spectrometry grade methanol. The methanol dilutions were mixed with a multichannel pipette set to 100 μL then centrifuged for 30 minutes at 3220 rcf. The centrifuged samples were then diluted 20X in ultrapure water containing 5.3 μM L-dopa-(*phenyl*-$d_3$) as an internal standard (10 μL in 190 μL) and mixed with a multichannel pipette before UPLC–MS/MS analysis.

**Inhibitor assays with purified TyrDC (Figures 4B, S7)**

TyrDC inhibitor assays followed a protocol similar to that reported previously, modified to allow increased depletion of the substrate[7].

The preincubation concentrations of each component were as follows: 1 μM TyrDC, 200 μM PLP, and 1-1000 μM inhibitor. The concentrations of each assay component were as follows: 0.1 μM TyrDC, 20 μM PLP, 0.1-100 μM inhibitor, and 500 μM L-dopa.

Inhibitors were dissolved to a concentration of 1.5 mM in TH buffer with vigorous shaking and sonication, followed by dilution in more TH buffer to give 300 μL aliquots of the desired concentrations. These aliquots were prepared one day prior to the assay and kept at 4 °C overnight. The next day, a 1.6 mM solution of PLP was prepared by dissolving 6.4 mg of PLP monohydrate in 15 mL of TH buffer. A 1 mM solution of L-dopa was made by dissolving 5.9 mg of L-dopa in 30 mL of NaOAc buffer, then 22.22 mL of the 1 mM solution was diluted with 17.78 mL of NaOAc buffer to give a 555 μM L-dopa solution. Inhibitor dilutions from the previous day were warmed to room temperature and distributed across triplicate PCR tubes in 30 μL aliquots. Reaction mixtures without inhibitor or without enzyme were included for comparison.

Flash-frozen TyrDC aliquots were thawed at 4 °C before warming to room temperature. The aliquots were diluted with TH buffer to yield an 8 μM TyrDC solution. The 8 μM TyrDC solution and 1.6 mM PLP solution were combined in a 1:1 ratio and mixed well using a micropipette to give a solution with 4 μM TyrDC and 800 μM PLP. Inhibitor preincubation was initiated by transferring 10 μL of the PLP/TyrDC mix to each inhibitor-containing PCR tube using a multichannel pipette and mixing well. After a 20 minute preincubation, the enzyme/inhibitor solutions were diluted 10X (20 μL in 180 μL) into the 555 μM L-dopa solution in 96-well plates. The reaction mixtures were mixed with a multichannel pipette set to 100 μL and left at room temperature for 20 additional minutes.

Reaction mixtures were mixed with a multichannel pipette set to 100 μL before being quenched by 10X dilution (20 μL in 180 μL) in mass spectrometry grade methanol. The methanol dilutions were mixed by multichannel pipette before centrifugation for 30 minutes at 3220 rcf. The centrifuged samples were then diluted 20X in ultrapure water containing 5.3 μM L-dopa-(*phenyl*-$d_3$) as an internal standard (10 μL in 190 μL) and mixed with a multichannel pipette before UPLC–MS/MS analysis.

**Inhibitor assays with purified AADC (Figure S8)**

AADC inhibitor assays followed a protocol similar to that reported previously, modified to allow increased depletion of the substrate[7].

The preincubation concentrations of each component were as follows: 2 μM AADC, 25 μM PLP, and 0.01-1000 μM inhibitor. The concentrations of each assay component were as follows: 0.2 μM AADC, 2.5 μM PLP, 0.001-100 μM inhibitor, and 250 μM L-dopa.

Inhibitors were dissolved to a concentration of 1.5 mM in TH buffer with vigorous shaking and sonication, followed by dilution in more TH buffer to give 300 μL aliquots of the desired concentrations. These aliquots were prepared one day prior to the assay and kept at 4 °C overnight. The next day, a 1 mM solution of PLP was prepared by dissolving 8 mg of PLP monohydrate in 30 mL of KPhos buffer, then 2.5 mL of the 1 mM solution was diluted with 22.5 mL of KPhos buffer to give a 100 μM PLP solution. A 1 mM solution of L-dopa was made by dissolving 5.9 mg of L-dopa in 30 mL of KPhos buffer, then 11.11 mL of the 1 mM solution was diluted with 28.89 mL of KPhos buffer to give a 278 μM L-dopa solution. Inhibitor dilutions from the previous day were warmed to room temperature and distributed across triplicate PCR tubes in 30 μL aliquots. Reaction mixtures without inhibitor or without enzyme were included for comparison.

Flash-frozen AADC aliquots were thawed at 4 °C before warming to room temperature. The aliquots were diluted with KPhos buffer containing 100 μM PLP to give a solution with 8 μM AADC and 100 μM PLP. Inhibitor preincubation was initiated by transferring 10 μL of the PLP/AADC mix to each inhibitor-containing PCR tube using a multichannel pipette and mixing well. After a 20 minute preincubation, the enzyme/inhibitor solutions were diluted 10X (20 μL in 180 μL) into the 278 μM L-dopa solution on 96-well plates. The reaction mixtures were mixed with a multichannel pipette set to 100 μL, after which the plates were sealed with foil plate seals and incubated at 37 °C for 30 additional minutes.

Reaction mixtures were mixed with a multichannel pipette set to 100 μL before being quenched by 10X dilution (20 μL in 180 μL) in mass spectrometry grade methanol. The methanol dilutions were mixed by multichannel pipette before centrifugation for 30 minutes at 3220 rcf. The centrifuged samples were then diluted 25X in ultrapure water containing 2.1 μM L-dopa-(*phenyl*-$d_3$) as an internal standard (8 μL in 192 μL) and mixed with a multichannel pipette before UPLC–MS/MS analysis.

**Inhibitor assays with a panel of enterococci (Figures 4C, S9)**

The concentration of L-dopa in assays was 500 μM and the concentration of inhibitors in assays was 25 μM. Inhibitors were dissolved to a concentration of 600 μM in BHI medium with vigorous shaking and sonication, after which the solutions were sterilized using 0.22 μm filters. 694 μL of sterilized inhibitor stock was diluted (using sterile technique) with 56 μL of BHI medium to provide 750 μL aliquots of 555 μM inhibitor. These aliquots were brought into an anaerobic chamber one day before use, as were the consumables needed for assay setup. A starter culture of each strain was inoculated from a glycerol stock in the anaerobic chamber using 8 mL of BHI medium in a Hungate tube. The tubes were sealed, brought out of the chamber, and cultured for 18 hours at 37 °C without shaking.

The next day, a 1 mM L-dopa solution was made in the chamber by dissolving 5.9 mg of L-dopa in 30 mL of BHI medium (this step requires sustained vortexing). 29.25 mL of the 1 mM L-dopa solution was diluted with 20.75 mL of BHI medium to give a 585 µM L-dopa solution. 9.5 mL of the 585 µM L-dopa solution was combined with 500 µL of the 555 µM inhibitor aliquots that had equilibrated overnight, providing solutions containing 555 µM L-dopa and 27.75 µM inhibitor. These solutions were mixed by vortexing, then distributed into 96-well plates in 180 µL aliquots. Starter cultures were diluted 10X into the inhibitor/L-dopa solutions (20 µL into 180 µL) using a multichannel pipette, after which the plates were sealed using foil plate seals and moved into a GasPak anaerobic pouch. Triplicate wells were included for each inhibitor/strain combination. The anaerobic pouch was removed from the chamber and placed in a 37 °C incubator for 18 hours without shaking.

After incubation, the plates were frozen at –80 °C for less than 5 hours before warming to room temperature and mixing with a multichannel pipette set to 100 µL. $OD_{600}$ was measured for all wells before 10X dilution (20 µL in 180 µL) in mass spectrometry grade methanol. The methanol dilutions were mixed with a multichannel pipette set to 100 µL then centrifuged for 30 minutes at 3220 rcf. The centrifuged samples were then diluted 20X in ultrapure water containing 5.3 µM L-dopa-(*phenyl*-$d_3$) as an internal standard (10 µL in 190 µL) and mixed with a multichannel pipette before UPLC–MS/MS analysis.

**Inhibitor assays with human fecal samples (Figures 4D, S10)**

Fecal samples from Parkinson's disease patients were obtained as part of the MICRO-PD clinical trial at the University of California San Francisco (ClinicalTrials.gov ID NCT03575195). The four samples used for inhibitor testing were collected at baseline, prior to any intervention, and stored at –80 °C until use. All four samples were provided by distinct donors.

The frozen fecal samples were brought into an anaerobic chamber and warmed to room temperature before suspending to a density of 0.1 g/mL in sterile phosphate-buffered saline. The resultant slurries were vortexed thoroughly to homogenize, after which they were left undisturbed for 45 minutes to allow particulates to settle. 200 µL slurry aliquots were combined with an equal volume of sterile 40% glycerol before being removed from the chamber, flash-frozen in liquid nitrogen, and stored at –80 °C until use. Blank aliquots were created following a similar workflow using phosphate-buffered saline without added fecal sample.

A flash-frozen aliquot of each fecal sample (and one blank) was warmed to room temperature in the anaerobic chamber and 60 µL of each aliquot was used to inoculate 6 mL of BHI medium in a Hungate tube. These starter cultures were sealed, brought out of the chamber, and grown for 48 hours at 37 °C without shaking.

The concentration of L-dopa in assays was 1000 μM and the concentration of inhibitors in assays was 250 μM. Inhibitors were dissolved to a concentration of 600 μM in BHI medium with vigorous shaking and sonication, after which the solutions were sterilized using 0.22 μm filters. Sterile inhibitor aliquots were brought into the anaerobic chamber one day before use, as were the consumables needed for assay setup.

The next day, a 2.5 mM L-dopa solution was made in the chamber by dissolving 14.8 mg of L-dopa in 30 mL of BHI medium (this step requires sustained vortexing). 17.46 mL of the 2.5 mM L-dopa solution was diluted with 2.54 mL of BHI medium to give a 2182 μM L-dopa solution. 1.048 mL of the 2182 μM L-dopa solution was combined with 0.952 mL of the 600 μM inhibitor aliquots that had equilibrated overnight, providing solutions with 1143 μM L-dopa and 285.6 μM inhibitor. These solutions were mixed by vortexing, then distributed into a 96-well plate in 113.75 μL aliquots. Starter cultures were diluted 8X into the inhibitor/L-dopa solutions (16.25 μL into 113.75 μL) using a multichannel pipette, after which the plate was sealed using a foil plate seal and moved into a GasPak anaerobic pouch. Triplicate wells were included for each inhibitor/fecal sample combination. The anaerobic pouch was removed from the chamber and placed in a 37 °C incubator for 40 hours without shaking.

After incubation, the plate was frozen at –80 °C for less than 3 hours before warming to room temperature. The undiluted samples were mixed with a multichannel pipette set to 100 μL before 20X dilution (10 μL in 190 μL) in mass spectrometry grade methanol. The methanol dilutions were mixed with a multichannel pipette set to 100 μL before centrifugation for 30 minutes at 3220 rcf. The centrifuged samples were then diluted 40X in ultrapure water containing 5.13 μM L-dopa-(*phenyl*-$d_3$) as an internal standard (5 μL in 195 μL) and mixed with a multichannel pipette before UPLC–MS/MS analysis.

### Synthetic methods

All inhibitors were obtained through custom synthesis services provided by WuXi AppTec. Enantiomers were separated as *tert*-butyl esters using preparative scale chiral SFC, and the active enantiomer was determined through inhibition assays with *E. faecalis* MMH594 (characterization data is only shown for the active enantiomer). The methods below are adapted directly from synthetic reports provided by WuXi AppTec.

*General scheme for the synthesis of* ***2*** *and* ***3***

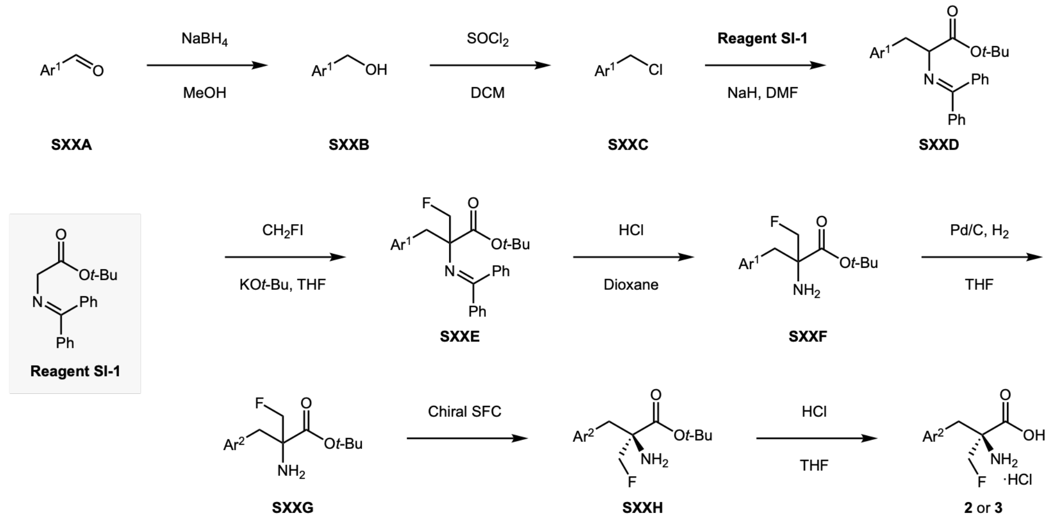


Here Ar[1] represents an arene with benzyl-protected hydroxylation, and Ar[2] represents the same arene after deprotection. XX represents 02 in the synthesis of **2** and 03 in the synthesis of **3**. Benzyl chloride **S02C** was purchased directly, so synthesis of **2** did not require the first two steps.

*General scheme for the synthesis of* ***26, 27,*** *and* ***28***

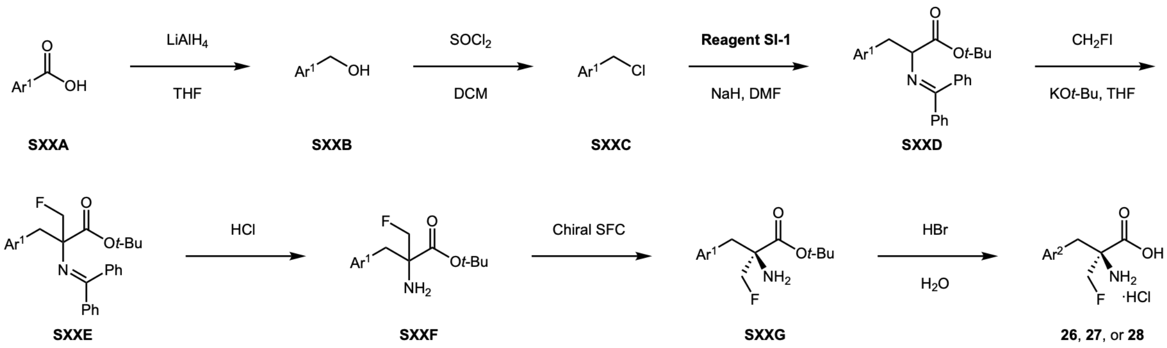


Here Ar[1] represents a difluoroarene with methyl-protected hydroxylation, and Ar[2] represents the same difluoroarene after deprotection. XX represents 26 in the synthesis of **26**, 27 in the synthesis of **27**, and 28 in the synthesis of **28**. Benzyl bromide **S28C** was purchased directly, so synthesis of **28** did not require the first two steps. Compound **S28G** was produced starting from **S28E** without isolation of the intermediate.

Synthesis of **2**

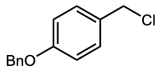

**S02C**

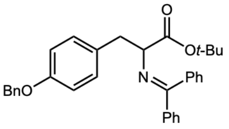

**S02D**

*Preparation of* ***S02D*** *from* ***S02C***

To a mixture of 1-benzyloxy-4-(chloromethyl)benzene (5.00 g, 21.49 mmol) and *tert*-butyl 2-(benzhydrylideneamino)acetate (5.8 g, 19.64 mmol) in DMF (80 mL) was added sodium hydride (863.90 mg, 21.60 mmol, 60% purity) in portions at 0 °C. After addition, the mixture was stirred at 20 °C for 2 h. After completion, the reaction mixture was quenched with sat. $NH_4Cl$ (200 mL) and extracted with EtOAc (100 mL x 3). The combined organic layers were dried over $Na_2SO_4$, filtered, and concentrated under vacuum. The residue was purified using column chromatography ($SiO_2$, petroleum ether/ethyl acetate = 50/1 to 20/1) to give *tert*-butyl 2-(benzhydrylideneamino)-3-(4-benzyloxyphenyl)propanoate (8.1 g, 16.48 mmol, 84% yield) as a yellow solid.

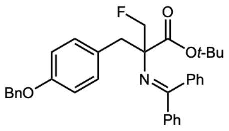

**S02E**

*Preparation of* ***S02E*** *from* ***S02D***

To a suspension of *tert*-butyl 2-(benzhydrylideneamino)-3-(4-benzyloxyphenyl)propanoate (8.1 g, 16.48 mmol) and fluoro(iodo)methane (9.22 g, 57.67 mmol) in THF (50 mL) was added dropwise KO*t*-Bu (57.67 mL of a 1 M solution in THF) at 20 °C. After addition, the mixture was stirred for 12 h. The reaction mixture was quenched with water (250 mL) and extracted with EtOAc (100 mL x 3). The combined organic layers were washed with brine (150 mL), dried over $Na_2SO_4$, filtered, and concentrated *in vacuo* to give *tert*-butyl 2-(benzhydrylideneamino)-2-[(4-benzyloxyphenyl)methyl]-3-fluoro-propanoate (7.9 g, crude) as a yellow oil that was used in the next step directly.

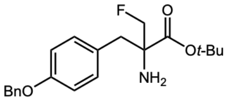

**S02F**

*Preparation of* ***S02F*** *from* ***S02E***

A mixture of *tert*-butyl 2-(benzhydrylideneamino)-2-[(4-benzyloxyphenyl)methyl]-3-fluoro-propanoate (7.9 g, 15.09 mmol) in HCl (0.5 M, 40.86 mL) and dioxane (40 mL) was stirred at 20 °C for 12 hours. The reaction mixture was quenched with sat. $NaHCO_3$ (200 mL) and extracted with EtOAc (100 mL x 3). The combined organic layers were washed with brine (200 mL), dried over $Na_2SO_4$, filtered, and concentrated *in vacuo*. The residue was purified using column chromatography ($SiO_2$, petroleum ether/ethyl acetate = 30/1 to 2/1) to give *tert*-butyl 2-amino-2-

[(4-benzyloxyphenyl)methyl]-3-fluoro-propanoate (4.8 g, 13.35 mmol, 89% yield) as a yellow solid.

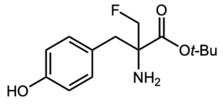


**S02G**

*Preparation of **S02G** from **S02F***

To a suspension of *tert*-butyl 2-amino-2-[(4-benzyloxyphenyl)methyl]-3-fluoro-propanoate (4.7 g, 13.08 mmol) in THF (200 mL) was added palladium, 10% on carbon (1 g, 13.08 mmol) (wetted with 55% water). The reaction mixture was stirred at 30 °C for 12 hours under $H_2$ (40 psi). After completion, the reaction mixture was filtered and washed with THF (300 mL). The filtrate was concentrated under vacuum. The residue was purified using column chromatography ($SiO_2$, ethyl acetate) to give *tert*-butyl 2-amino-2-(fluoromethyl)-3-(4-hydroxyphenyl)propanoate (2.8 g, 10.40 mmol, 80% yield) as a light yellow solid.

$^1$H NMR (400 MHz, MeOD) δ ppm 7.04 (d, $J$ = 8.2 Hz, 2H), 6.73 (d, $J$ = 8.4 Hz, 2H), 4.74-4.60 (m, 1H), 4.43-4.29 (m, 1H), 2.95 (d, $J$ = 13.6 Hz, 1H), 2.67 (d, $J$ = 13.6 Hz, 1H), 1.47 (s, 9H)

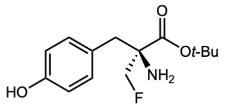


**S02H**

*Separation of **S02H** from **S02G***

*Tert*-butyl 2-amino-2-(fluoromethyl)-3-(4-hydroxyphenyl)propanoate (2.8 g, 10.40 mmol) was separated using chiral SFC (instrument: Waters SFC150AP preparative SFC; column: Daicel CHIRALPAK AD (250 x 30 mm, 10 µm); mobile phase: A $CO_2$ and B EtOH (0.1% $NH_3H_2O$); gradient: B% = 18% isocratic elution mode; flow rate: 60 g/min; wavelength: 220 nm; column temperature: 35 °C; system back pressure: 100 bar) to give *tert*-butyl (2*S*)-2-amino-2-(fluoromethyl)-3-(4-hydroxyphenyl)propanoate (1.3 g, 4.83 mmol, 46% yield) as a light yellow oil.

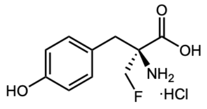


**2**

*Preparation of **2** from **S02H***

A mixture of *tert*-butyl (2*S*)-2-amino-2-(fluoromethyl)-3-(4-hydroxyphenyl)propanoate (1.29 g, 4.79 mmol) in 6 N HCl (6 M, 14.88 mL) and THF (15 mL) was stirred at 70 °C for 12 h. After completion, the reaction mixture was concentrated under vacuum. The residue was combined with that of a 100 mg scale reaction and purified using preparative HPLC (column: Phenomenex Luna C18 (250 x 70 mm, 15 µm); mobile phase: A water (0.04% HCl) and B ACN; gradient: B% = 1%-

100% over 20 min) to give 922 mg of (2*S*)-2-amino-2-(fluoromethyl)-3-(4-hydroxyphenyl)propanoic acid (99.93% purity, HCl salt) as a white solid.

$^1$H NMR (400 MHz, DMSO-$d_6$) δ ppm 9.52 (br s, 1H), 8.66 (br s, 2H), 7.03 (d, $J$ = 8.38 Hz, 2H), 6.71 (d, $J$ = 8.38 Hz, 2H), 4.90-4.76 (m, 1H), 4.71-4.56 (m, 1H), 2.95-3.05 (m, 2H)

Synthesis of **3**

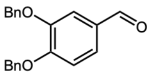


**S03A**

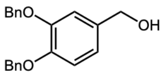


**S03B**

*Preparation of* ***S03B*** *from* ***S03A***

To a suspension of 3,4-dibenzyloxybenzaldehyde (12 g, 37.69 mmol) in methanol (128.42 mL), $NaBH_4$ (25.38 g, 58.15 mmol) was added in portions at 0 °C and the reaction mixture was stirred at 20 °C for 12 hours. After completion, the reaction mixture was concentrated under vacuum. The residue was acidified with 1 N HCl (200 mL) and extracted with EtOAc (100 mL x 3). The combined organic layers were washed with sat. $NaHCO_3$ (200 mL), washed with brine (200 mL), dried over $Na_2SO_4$, filtered, and concentrated *in vacuo* to give (3,4-dibenzyloxyphenyl)methanol (12 g, 37.46 mmol, 99% yield) as a brown solid. It was used in the next step directly without purification.

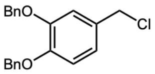


**S03C**

*Preparation of* ***S03C*** *from* ***S03B***

To a suspension of (3,4-dibenzyloxyphenyl)methanol (12 g, 37.46 mmol) in DCM (150 mL), thionyl chloride (6.68 g, 56.18 mmol) was added dropwise at 0 °C and the reaction mixture was stirred at 20 °C for 12 hours. After completion, the reaction mixture was concentrated under vacuum. The residue was purified using column chromatography ($SiO_2$, petroleum ether/ethyl acetate = 5/1) to give 1,2-dibenzyloxy-4-(chloromethyl)benzene (12 g, 35.42 mmol, 95% yield) as a light yellow solid.

$^1H$ NMR (400 MHz, $CDCl_3$) δ ppm 7.53 - 7.30 (m, 10H), 7.01 (s, 1H), 6.91 (s, 2H), 5.18 (d, $J$ = 2.1 Hz, 4H), 4.52 (s, 2H)

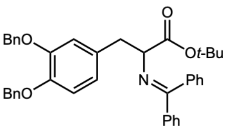


**S03D**

*Preparation of* ***S03D*** *from* ***S03C***

To a suspension of *tert*-butyl 2-(benzhydrylideneamino)acetate (4.8 g, 16.25 mmol) and 1,2-dibenzyloxy-4-(chloromethyl)benzene (6.06 g, 17.88 mmol) in DMF (80 mL) was added sodium hydride (715.02 mg, 17.88 mmol, 60% purity) in portions at 0 °C. The mixture was stirred at 20 °C for 2 h. After completion, the reaction mixture was quenched with ice water (200 mL) and extracted with EtOAc (100 mL x 3). The combined organic layers were washed with brine (200 mL), dried over $Na_2SO_4$, filtered, and concentrated under vacuum. The residue was purified using column chromatography ($SiO_2$, petroleum ether/ethyl acetate = 50/1 to 20/1) to give *tert*-butyl 2-

(benzhydrylideneamino)-3-(3,4-dibenzyloxyphenyl)propanoate (7.3 g, 12.21 mmol, 75% yield) as a yellow oil.

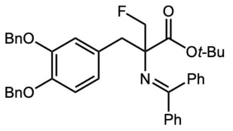


**S03E**

*Preparation of **S03E** from **S03D***

To a suspension of *tert*-butyl 2-(benzhydrylideneamino)-3-(3,4-dibenzyloxyphenyl)propanoate (7.3 g, 12.21 mmol) and fluoro(iodo)methane (7.03 g, 43.97 mmol) in THF (40 mL) was added KO*t*-Bu (43.97 mL of a 1 M solution in THF) dropwise and the reaction mixture was stirred at 20 °C for 12 hours. After completion, the reaction mixture was quenched with water (200 mL) and extracted with EtOAc (100 mL x 3). The combined organic layers were washed with brine (150 mL), dried over $Na_2SO_4$, filtered, and concentrated *in vacuo* to give *tert*-butyl 2-(benzhydrylideneamino)-2-[(3,4-dibenzyloxyphenyl)methyl]-3-fluoro-propanoate (7.6 g, 12.07 mmol, 99% yield) as a yellow oil. It was used in the next step without purification.

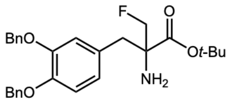


**S03F**

*Preparation of **S03F** from **S03E***

A mixture of *tert*-butyl 2-(benzhydrylideneamino)-2-[(3,4-dibenzyloxyphenyl)methyl]-3-fluoro-propanoate (7.6 g, 12.07 mmol) in 0.5 N HCl (0.5 M, 41.64 mL) and dioxane (41 mL) was stirred at 20 °C for 12 hours. The reaction mixture was quenched with sat. $NaHCO_3$ (100 mL) and extracted with EtOAc (50 mL x 3). The combined organic layers were washed with brine (100 mL), dried over $Na_2SO_4$, filtered, and concentrated under vacuum. The residue was purified using column chromatography ($SiO_2$, petroleum ether/ethyl acetate = 15/1 to 2/1) to give *tert*-butyl 2-amino-2-[(3,4-dibenzyloxyphenyl)methyl]-3-fluoro-propanoate (4 g, 8.59 mmol, 71% yield) as a yellow oil.

$^1$H NMR (400 MHz, $CDCl_3$) δ ppm 7.45 - 7.31 (m, 10H), 6.86 (d, $J$ = 8.2 Hz, 1H), 6.82 (d, $J$ = 2.0 Hz, 1H), 6.71 (dd, $J$ = 2.0, 8.1 Hz, 1H), 5.15 (s, 4H), 4.71 - 4.46 (m, 1H), 4.38 - 4.15 (m, 1H), 2.92 (d, $J$ = 13.3 Hz, 1H), 2.64 (d, $J$ = 13.3 Hz, 1H), 1.44 (s, 9H)

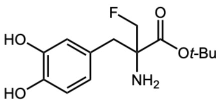


**S03G**

*Preparation of **S03G** from **S03F***

To a suspension of *tert*-butyl 2-amino-2-[(3,4-dibenzyloxyphenyl)methyl]-3-fluoro-propanoate (3.9 g, 8.38 mmol) in THF (100 mL) was added palladium, 10% on carbon (2 g, 8.38 mmol) (wetted with 55% water). The reaction mixture was stirred at 20 °C for 12 hours under $H_2$ (40 psi). After completion, the reaction mixture was filtered and washed with MeOH (100 mL). The filtrate was concentrated under vacuum. The residue was triturated with DCM (15 mL) and filtered, then the filter cake was dried *in vacuo* to give *tert*-butyl 2-amino-2-[(3,4-dihydroxyphenyl)methyl]-3-fluoro-propanoate (1.2 g, 4.21 mmol, 50% yield) as a white solid.

$^{1}$H NMR (400 MHz, MeOD) δ ppm 6.71 (d, *J* = 8.1 Hz, 1H), 6.66 (d, *J* = 2.1 Hz, 1H), 6.53 (dd, *J* = 2.1, 8.1 Hz, 1H), 4.76 - 4.55 (m, 1H), 4.48 - 4.26 (m, 1H), 2.90 (d, *J* = 13.4 Hz, 1H), 2.60 (d, *J* = 13.4 Hz, 1H), 1.47 (s, 9H)

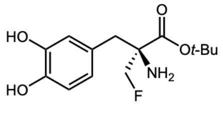


**S03H**

*Separation of **S03H** from **S03G***

*Tert*-butyl 2-amino-2-[(3,4-dihydroxyphenyl)methyl]-3-fluoro-propanoate (500 mg, 1.75 mmol) was separated using chiral SFC (instrument: Thar SFC80 preparative SFC; Column: Daicel CHIRALPAK AD (250 x 30 mm, 10 µm); mobile phase: A $CO_2$ and B MeOH (0.1% $NH_3H_2O$); gradient: B% = 30% isocratic elution mode; flow rate: 55 g/min; wavelength: 220 nm; column temperature: 40 °C; system back pressure: 100 bar) to give *tert*-butyl (2*S*)-2-amino-2-[(3,4-dihydroxyphenyl)methyl]-3-fluoro-propanoate (200 mg, 695.80 µmol, 40% yield, 99.26% purity) as a light yellow oil.

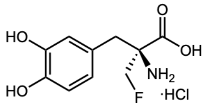


**3**

*Preparation of **3** from **S03H***

A mixture of *tert*-butyl (2*S*)-2-amino-2-[(3,4-dihydroxyphenyl)methyl]-3-fluoro-propanoate (150.00 mg, 525.74 µmol) in 6 N HCl (6 M, 1.50 mL) and THF (1.5 mL) was stirred at 70 °C for 12 h. After completion, the reaction mixture was concentrated under vacuum. The residue was combined with that of a 50 mg scale reaction and purified using preparative HPLC (column: Welch Xtimate C18 (100 x 25 mm, 3 µm); mobile phase: A water (0.04% HCl) and B ACN; gradient: B% = 1% for 8 min) to give 130.1 mg of (2*S*)-2-amino-2-[(3,4-dihydroxyphenyl)methyl]-3-fluoro-propanoic acid (99.86% purity, HCl salt) as a white solid.

$^{1}$H NMR (400 MHz, DMSO-$d_6$) δ ppm 8.89 (br d, *J* = 18.2 Hz, 3H), 6.66 - 6.63 (m, 2H), 6.48 (dd, *J* = 2.1, 8.1 Hz, 1H), 4.84 - 4.69 (m, 1H), 4.63 - 4.49 (m, 1H), 2.93 (d, *J* = 14 Hz, 1H), 2.83 (d, *J* = 14 Hz, 1H)

Synthesis of **26**

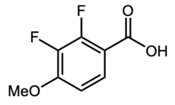

**S26A**

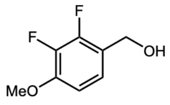

**S26B**

*Preparation of **S26B** from **S26A***

To a solution of 2,3-difluoro-4-methoxy-benzoic acid (2.5 g, 13.29 mmol) in THF (25 mL) was added $LiAlH_4$ (756.55 mg, 19.93 mmol) slowly at 0 °C. The mixture was stirred at 20 °C for 5 hr. The reaction mixture was quenched with aq. $NH_4Cl$ (100 mL) at 0 °C, then diluted with $H_2O$ (50 mL) and extracted with EtOAc (50 mL x 2). The combined organic layers were concentrated under reduced pressure to give a residue. The residue was purified using column chromatography ($SiO_2$, petroleum ether/ethyl acetate = 20/1 to 1/1) to give (2,3-difluoro-4-methoxy-phenyl)methanol (1.8 g, 10.34 mmol, 78% yield) as a white solid.

$^1$H NMR (400 MHz, $CDCl_3$) δ ppm 3.91 (s, 3H), 4.70 (s, 2H), 6.69 - 6.80 (m, 1H), 7.03 - 7.13 (m, 1H)

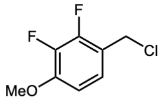

**S26C**

*Preparation of **S26C** from **S26B***

To a solution of (2,3-difluoro-4-methoxy-phenyl)methanol (1.8 g, 10.34 mmol) in DCM (18 mL) was added $SOCl_2$ (3.69 g, 31.01 mmol, 2.25 mL) at 0 °C. The mixture was stirred at 20 °C for 5 hr. The reaction mixture was diluted with aq. $NaHCO_3$ (50 mL) and extracted with DCM (50 mL x 2). The combined organic layers were concentrated under reduced pressure to give 1-(chloromethyl)-2,3-difluoro-4-methoxy-benzene (1.4 g, 7.27 mmol, 70% yield) as a colorless oil. The residue was used in the next step without further purification.

$^1$H NMR (400 MHz, $CDCl_3$) δ ppm 3.92 (s, 3H), 4.61 (d, *J* = 0.88 Hz, 2H), 6.74 (ddd, *J* = 8.88, 7.34, 1.97 Hz, 1H), 7.10 (td, *J* = 8.22, 2.41 Hz, 1H)

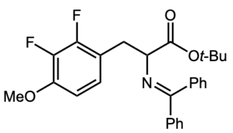

**S26D**

*Preparation of **S26D** from **S26C***

To a solution of 1-(chloromethyl)-2,3-difluoro-4-methoxy-benzene (1.4 g, 7.27 mmol) and *tert*-butyl 2-(benzhydrylideneamino)acetate (1.95 g, 6.61 mmol) in DMF (14 mL) was added NaH (290.75 mg, 7.27 mmol, 60% purity) slowly at 0 °C. The mixture was stirred at 25 °C for 2 hr under $N_2$. The reaction mixture was quenched by addition of aq. $NH_4Cl$ (50 mL) at 0 °C, then diluted with $H_2O$ (50 mL) and extracted with DCM (50 mL x 2). The combined organic layers were concentrated under reduced pressure to give a residue. The residue was purified using column chromatography ($SiO_2$, petroleum ether/ethyl acetate = 20/1 to 3/1), giving *tert*-butyl 2-(benzhydrylideneamino)-3-(2,3-difluoro-4-methoxy-phenyl)propanoate (2.1 g, 4.65 mmol, 70% yield) as a yellow solid.

$^1$H NMR (400 MHz, $CDCl_3$) δ ppm 1.45 (s, 9H), 3.27 (br d, $J$ = 3.91 Hz, 2H), 3.86 (s, 3H), 4.14 - 4.25 (m, 1H), 6.53 - 6.63 (m, 1H), 6.78 (br d, $J$ = 6.85 Hz, 2H), 6.81 - 6.91 (m, 1H), 7.29 - 7.44 (m, 6H), 7.60 (br d, $J$ = 7.82 Hz, 2H)

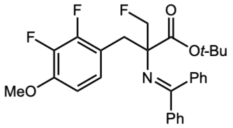


**S26E**

*Preparation of* ***S26E*** *from* ***S26D***

To a solution of *tert*-butyl 2-(benzhydrylideneamino)-3-(2,3-difluoro-4-methoxy-phenyl)propanoate (2.1 g, 4.65 mmol) and fluoro(iodo)methane (2.60 g, 16.28 mmol) in THF (21 mL) was added KO*t*-Bu (16.28 mL of a 1 M solution in THF) slowly at 25 °C. The mixture was stirred at 25 °C for 12 hr. The reaction mixture was diluted with $H_2O$ (200 mL) and extracted with EtOAc (100 mL x 2). The combined organic layers were washed with brine (150 mL), dried over $Na_2SO_4$, filtered, and concentrated under reduced pressure to give a residue, which was used in the next step without further purification. *Tert*-butyl 2-(benzhydrylideneamino)-2-[(2,3-difluoro-4-methoxy-phenyl)methyl]-3-fluoro-propanoate (1.2 g, 2.48 mmol, 53% yield) was obtained as a white solid.

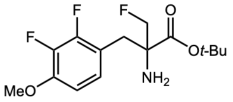


**S26F**

*Preparation of* ***S26F*** *from* ***S26E***

A solution of *tert*-butyl 2-(benzhydrylideneamino)-2-[(2,3-difluoro-4-methoxy-phenyl)methyl]-3-fluoro-propanoate (1.2 g, 2.48 mmol) in HCl (0.5 M, 6 mL) and dioxane (6 mL) was stirred at 20 °C for 12 hr under $N_2$. The reaction mixture was diluted with aq. $NaHCO_3$ (200 mL) at 0 °C and extracted with EtOAc (100 mL x 2). The combined organic layers were washed with brine (200 mL), dried over $Na_2SO_4$, filtered, and concentrated under reduced pressure to give a residue. The residue was purified using column chromatography ($SiO_2$, petroleum ether/ethyl acetate = 20/1 to 0/1). The residue was purified using preparative HPLC (column: Phenomenex C18 (75 x 30 mm, 3 µm); mobile phase: A water ($NH_3H_2O$, $NH_4HCO_3$) and B ACN; gradient: B% = 35%-70% over 8 min). *Tert*-butyl 2-amino-2-[(2,3-difluoro-4-methoxy-phenyl)methyl]-3-fluoro-propanoate (0.53 g, 1.66 mmol, 67% yield) was obtained as a yellow oil.

$^1$H NMR (400 MHz, $CDCl_3$) δ ppm 1.46 (s, 9H), 2.79 - 2.91 (m, 1H), 2.97 - 3.05 (m, 1H), 3.90 (s, 3H), 4.31 (d, *J* = 8.58 Hz, 0.5H), 4.43 (d, *J* = 8.58 Hz, 0.5H), 4.61 (s, 0.5H), 4.74 (d, *J* = 8.70 Hz, 0.5H), 6.70 (s, 1H), 6.92 - 6.99 (m, 1H)

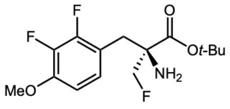


**S26G**

*Separation of **S26G** from **S26F***

*Tert*-butyl 2-amino-2-(2,3-difluoro-4-methoxybenzyl)-3-fluoropropanoate (0.53 g, 1.66 mmol) was separated using chiral SFC (column: Daicel CHIRALPAK IC (250 x 30 mm, 10 µm); gradient: B% = 10% for 12 min). *Tert*-butyl (2*S*)-2-amino-2-[(2,3-difluoro-4-methoxy-phenyl)methyl]-3-fluoro-propanoate (140 mg, 438.43 µmol, 26% yield) was obtained as a yellow oil.

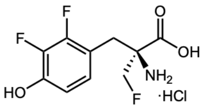


**26**

*Preparation of **26** from **S26G***

A solution of *tert*-butyl (2*S*)-2-amino-2-[(2,3-difluoro-4-methoxy-phenyl)methyl]-3-fluoro-propanoate (130 mg, 407.12 µmol) in HBr (68.63 mg, 407.12 µmol, 0.5 mL, 48% purity) was stirred at 110 °C for 12 hr under $N_2$. The reaction mixture was concentrated under reduced pressure to give a residue. The residue was purified using preparative HPLC (column: Phenomenex Luna C18 (80 x 40 mm, 3 µm); mobile phase: A water (HCl) and B ACN; gradient: B% = 5%-20% over 7 min) to give (2*S*)-2-amino-2-[(2,3-difluoro-4-hydroxy-phenyl)methyl]-3-fluoro-propanoic acid (60 mg, 240.78 µmol, 59% yield, HCl salt) as a white solid.

$^1$H NMR (400 MHz, DMSO-$d_6$) δ ppm 2.98 (s, 2H), 4.48 (d, *J* = 10.01 Hz, 0.5H), 4.62 (dd, *J* = 17.76, 10.01 Hz, 1H), 4.76 (d, *J* = 9.88 Hz, 0.5H), 6.65 - 6.78 (m, 1H), 6.89 (td, *J* = 8.32, 1.75 Hz, 1H), 7.49 - 8.86 (m, 2H), 10.45 (br s, 1H)

Synthesis of **27**

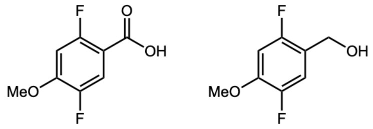


**S27A** **S27B**

*Preparation of* ***S27B*** *from* ***S27A***

To a solution of 2,5-difluoro-4-methoxy-benzoic acid (2.5 g, 13.29 mmol) in THF (25 mL) was added $LiAlH_4$ (756.55 mg, 19.93 mmol) slowly at 0 °C. The mixture was stirred at 20 °C for 5 hr. The reaction mixture was quenched with aq. $NH_4Cl$ (50 mL) at 0 °C, then diluted with $H_2O$ (50 mL) and extracted with EtOAc (50 mL x 2). The combined organic layers were concentrated under reduced pressure to give a residue. The residue was purified using column chromatography ($SiO_2$, petroleum ether/ethyl acetate = 20/1 to 1/1), giving (2,5-difluoro-4-methoxy-phenyl)methanol (1.7 g, 9.76 mmol, 73% yield) as a white solid.

$^1$H NMR (400 MHz, $CDCl_3$) δ ppm 1.89 (s, 1H), 3.87 (s, 3H), 4.66 (s, 2H), 6.69 (dd, *J* = 10.96, 7.02 Hz, 1H), 7.14 (dd, *J* = 11.18, 7.02 Hz, 1H)

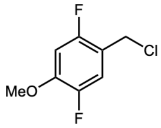


**S27C**

*Preparation of* ***S27C*** *from* ***S27B***

To a solution of (2,5-difluoro-4-methoxy-phenyl)methanol (1.7 g, 9.76 mmol) in DCM (20 mL) was added $SOCl_2$ (3.48 g, 29.29 mmol, 2.12 mL). The mixture was stirred at 25 °C for 5 hr. The reaction mixture was diluted with aq. $NaHCO_3$ (20 mL) and extracted with DCM (10 mL x 2). The combined organic layers were concentrated under reduced pressure to give 1-(chloromethyl)-2,5-difluoro-4-methoxy-benzene (1.5 g, 7.79 mmol, 80% yield) as a colorless oil. The residue was used in the next step without further purification.

$^1$H NMR (400 MHz, $CDCl_3$) δ ppm 3.89 (s, 3H), 4.57 (s, 2H), 6.71 (dd, *J* = 10.85, 6.91 Hz, 1H), 7.13 (dd, *J* = 11.07, 6.91 Hz, 1H)

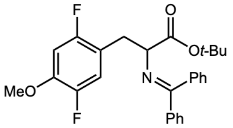


**S27D**

*Preparation of* ***S27D*** *from* ***S27C***

To a solution of 1-(chloromethyl)-2,5-difluoro-4-methoxy-benzene (1.5 g, 7.79 mmol) and *tert*-butyl 2-(benzhydrylideneamino)acetate (2.09 g, 7.08 mmol) in DMF (15 mL) was added NaH (311.51 mg, 7.79 mmol, 60% purity) slowly at 0 °C. The mixture was stirred at 25 °C for 2 hr under $N_2$. The reaction mixture was quenched by addition of aq. $NH_4Cl$ (100 mL) at 0 °C, then diluted with $H_2O$ (200 mL) and extracted with DCM (200 mL x 2). The combined organic layers were concentrated under reduced pressure to give a residue. The residue was purified using column chromatography ($SiO_2$, petroleum ether/ethyl acetate = 20/1 to 3/1), giving *tert*-butyl 2-(benzhydrylideneamino)-3-(2,5-difluoro-4-methoxy-phenyl)propanoate (2 g, 4.43 mmol, 63% yield) as a yellow solid.

$^1$H NMR (400 MHz, $CDCl_3$) δ ppm 1.44 (s, 9H), 2.99 - 3.26 (m, 2H), 3.82 (s, 3H), 4.15 (br s, 1H), 6.57 (dd, *J* = 10.58, 7.06 Hz, 1H), 6.82 (br d, *J* = 7.28 Hz, 2H), 6.84 - 6.91 (m, 1H), 7.28 - 7.39 (m, 6H), 7.58 (br d, *J* = 7.50 Hz, 2H)

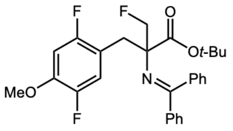


**S27E**

*Preparation of* ***S27E*** *from* ***S27D***

To a solution of *tert*-butyl 2-(benzhydrylideneamino)-3-(2,5-difluoro-4-methoxy-phenyl)propanoate (2 g, 4.43 mmol) and fluoro(iodo)methane (2.48 g, 15.50 mmol) in THF (20 mL) was added KO*t*-Bu (15.50 mL of a 1 M solution in THF) slowly at 25 °C. The mixture was stirred at 25 °C for 12 hr under $N_2$. The reaction mixture was diluted with $H_2O$ (50 mL) and extracted with EtOAc (50 mL x 2). The combined organic layers were washed with brine (150 mL), dried over $Na_2SO_4$, filtered, and concentrated under reduced pressure to give *tert*-butyl 2-(benzhydrylideneamino)-2-[(2,5-difluoro-4-methoxy-phenyl)methyl]-3-fluoro-propanoate (1.8 g, 3.72 mmol, 84% yield) as a white solid. The residue was used in the next step without further purification.

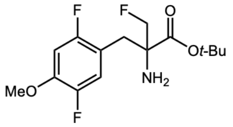


**S27F**

*Preparation of* ***S27F*** *from* ***S27E***

A solution of *tert*-butyl 2-(benzhydrylideneamino)-2-[(2,5-difluoro-4-methoxy-phenyl)methyl]-3-fluoro-propanoate (1.8 g, 3.72 mmol) in HCl (0.5 M, 9 mL) and dioxane (9 mL) was stirred at 20 °C for 12 hr under $N_2$. The reaction mixture was diluted with aq. $NaHCO_3$ (200 mL) at 0 °C and extracted with EtOAc (100 mL x 2). The combined organic layers were washed with brine (200 mL), dried over $Na_2SO_4$, filtered, and concentrated under reduced pressure to give a residue which was purified using column chromatography ($SiO_2$, petroleum ether/ethyl acetate = 20/1 to 0/1). The residue was purified using preparative HPLC (column: Waters Xbridge BEH C18 (250 x 50 mm, 10 μm); mobile phase: A water ($NH_3H_2O$, $NH_4HCO_3$) and B ACN; gradient: B% = 35%-65% over 10 min). *Tert*-butyl 2-amino-2-[(2,5-difluoro-4-methoxy-phenyl)methyl]-3-fluoro-propanoate (0.56 g, 1.75 mmol, 47% yield) was obtained as a yellow oil.

$^1$H NMR (400 MHz, $CDCl_3$) δ ppm 1.47 (s, 9H), 2.78 - 2.85 (m, 1H), 2.91 - 2.97 (m, 1H), 3.87 (s, 3H), 4.29 (d, $J$ = 8.58 Hz, 0.5H), 4.41 (d, $J$ = 8.58 Hz, 0.5H), 4.60 (d, $J$ = 8.70 Hz, 0.5H), 4.71 (d, $J$ = 8.58 Hz, 0.5H), 6.69 (dd, $J$ = 10.73, 7.15 Hz, 1H), 6.99 (dd, $J$ = 11.56, 6.91 Hz, 1H)

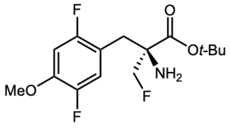


**S27G**

*Separation of **S27G** from **S27F***

*Tert*-butyl 2-amino-2-(2,5-difluoro-4-methoxybenzyl)-3-fluoropropanoate (0.56 g, 1.75 mmol) was separated using chiral SFC (column: Daicel CHIRALPAK IC (250 x 30 mm, 10 μm); gradient: B% = 10% for 12 min). *Tert*-butyl (2*S*)-2-amino-2-[(2,5-difluoro-4-methoxy-phenyl)methyl]-3-fluoro-propanoate (140 mg, 438.43 μmol, 25% yield) was obtained as a yellow oil.

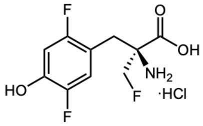


**27**

*Preparation of **27** from **S27G***

A solution of *tert*-butyl (2*S*)-2-amino-2-[(2,5-difluoro-4-methoxy-phenyl)methyl]-3-fluoro-propanoate (130 mg, 407.12 μmol) in HBr (68.63 mg, 407.12 μmol, 0.5 mL, 48% purity) was stirred at 110 °C for 12 hr under $N_2$. The reaction mixture was concentrated under reduced pressure to give a residue. The residue was purified using preparative HPLC (column: Phenomenex Luna C18 (80 x 40 mm, 3 μm); mobile phase: A water (HCl) and B ACN; gradient: B% = 5%-20% over 7 min) to give (2*S*)-2-amino-2-[(2,5-difluoro-4-hydroxy-phenyl)methyl]-3-fluoro-propanoic acid (60 mg, 240.78 μmol, 59% yield, HCl salt) as a white solid.

$^1$H NMR (400 MHz, DMSO-$d_6$) δ ppm 2.86 - 2.98 (m, 2H), 4.42 (d, $J$ = 9.76 Hz, 0.5H), 4.54 (d, $J$ = 9.88 Hz, 0.5H), 4.61 (d, $J$ = 9.76 Hz, 0.5H), 4.73 (d, $J$ = 9.88 Hz, 0.5H), 6.73 (dd, $J$ = 10.76, 7.50 Hz, 1H), 7.09 (dd, $J$ = 11.69, 7.19 Hz, 1H), 8.14 (br s, 2H), 10.25 - 10.71 (m, 1H)

Synthesis of **28**

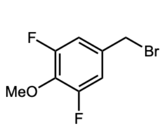

**S28C**

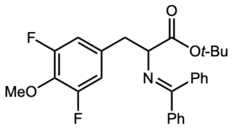

**S28D**

*Preparation of **S28D** from **S28C***

To a solution of 5-(bromomethyl)-1,3-difluoro-2-methoxy-benzene (5.5 g, 23.20 mmol) and *tert*-butyl 2-(benzhydrylideneamino)acetate (6.85 g, 23.20 mmol) in DMF (50 mL) was added NaH (1.11 g, 27.84 mmol, 60% purity) at 0 °C under $N_2$ atmosphere. The mixture was stirred at 20 °C for 8 hrs. The reaction mixture was quenched by addition of $H_2O$ (20 mL) at 0 °C under $N_2$ atmosphere. The mixture was extracted with EtOAc (10 mL x 3), then the combined organic layers were dried over $Na_2SO_4$, filtered, and concentrated under reduced pressure to give a residue. The residue was purified using column chromatography ($SiO_2$, petroleum ether/ethyl acetate = 1/0 to 10/1) to give *tert*-butyl 2-(benzhydrylideneamino)-3-(3,5-difluoro-4-methoxy-phenyl)propanoate (5.9 g, 11.24 mmol, 48% yield, 86% purity) as a white solid.

$^1$H NMR (400 MHz, $CDCl_3$) δ ppm 7.81 (d, $J$ = 7.2 Hz, 1H), 7.55 (d, $J$ = 1.2 Hz, 2H), 7.35 - 7.41 (m, 6H), 6.81 (d, $J$ = 4.1 Hz, 1H), 6.63 (d, $J$ = 8 Hz, 2H), 4.11 - 4.37 (m, 1H), 3.96 (s, 3H), 3.07 - 3.23 (m, 2H), 1.43 (s, 9H)

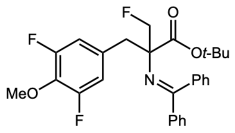

**S28E**

*Preparation of **S28E** from **S28D***

To a solution of *tert*-butyl 2-(benzhydrylideneamino)-3-(3,5-difluoro-4-methoxy-phenyl)propanoate (2.88 g, 6.38 mmol) in THF (30 mL) was added fluoro(iodo)methane (5.1 g, 31.89 mmol) and KO*t*-Bu (3.58 g, 31.89 mmol). The mixture was stirred at 20 °C for 8 hrs. The mixture was diluted with EtOAc (40 mL), washed with water (20 mL x 3), washed with brine (20 mL x 2), dried with anhydrous $Na_2SO_4$, filtered, and concentrated under reduced pressure to give a residue. The residue was purified using column chromatography ($SiO_2$, petroleum ether/ethyl acetate = 1/0 to 10/1) to give *tert*-butyl 2-(benzhydrylideneamino)-2-[(3,5-difluoro-4-methoxy-phenyl)methyl]-3-fluoro-propanoate (1.4 g, 2.78 mmol, 44% yield, 96% purity) as a white solid.

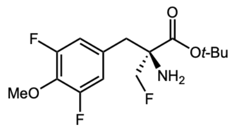

**S28G**

*Preparation of **S28G** from **S28E***

To a solution of *tert*-butyl 2-(benzhydrylideneamino)-2-[(3,5-difluoro-4-methoxy-phenyl)methyl]-3-fluoro-propanoate (0.8 g, 1.65 mmol) in THF (10 mL) was added HCl (1 M, 1.65 mL). The mixture was stirred at 20 °C for 1 hr. The reaction mixture was diluted with $H_2O$ (2 mL) and washed with petroleum ether (2 mL x 3). The crude mixture was adjusted to pH 7 with saturated aqueous $NaHCO_3$ and extracted with EtOAc (5 mL x 3). The combined organic layers were dried over $Na_2SO_4$, filtered, and concentrated under reduced pressure to give a residue. The residue was separated using chiral SFC (column: Daicel CHIRALPAK AD (250 x 30 mm, 10 µm); mobile phase: A $CO_2$ and B 1:1 Hep, EtOH (0.1% $NH_3H_2O$); gradient: B% = 11% isocratic elution mode) to give *tert*-butyl (2*S*)-2-amino-2-[(3,5-difluoro-4-methoxy-phenyl)methyl]-3-fluoro-propanoate (190 mg, 541.47 µmol, 33% yield, 91% purity) as a white solid, 99% purity by SFC. NMR analysis was conducted prior to chiral separation.

$^1$H NMR (400 MHz, $CDCl_3$) δ ppm 6.78-6.80 (m, 2H), 4.30-4.69 (m, 2H), 3.98 (s, 3H), 2.95 (d, *J* = 13.2 Hz, 1H), 2.67 (d, *J* = 13.2 Hz, 1H), 1.48 (s, 9H)

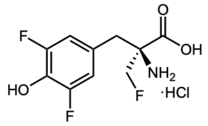


**28**

*Preparation of **28** from **S28G***

A solution of *tert*-butyl (2*S*)-2-amino-2-[(3,5-difluoro-4-methoxy-phenyl)methyl]-3-fluoro-propanoate (327 mg, 1.02 mmol) in HBr (5 mL, 40%) was degassed and purged with $N_2$ three times, then the mixture was stirred at 100 °C for 5 hrs under $N_2$ atmosphere. The crude reaction mixture was combined with another batch and adjusted to pH 7 with saturated aqueous $NaHCO_3$ before concentration under reduced pressure to get a residue. The residue was purified using preparative HPLC (column: Phenomenex Luna C18 (75 x 30 mm, 3 µm); mobile phase: A $H_2O$ (0.04% HCl) and B ACN; gradient: B% = 1%-8% over 8 min) to give (2*S*)-2-amino-2-[(3,5-difluoro-4-hydroxy-phenyl)methyl]-3-fluoro-propanoic acid (140 mg, 485.21 µmol, 41% yield, 99% purity, HCl salt) as a white solid.

$^1$H NMR (400 MHz, DMSO-$d_6$) δ ppm 10.27 (s, 1H), 8.70 (s, 2H), 6.91 (d, *J* = 8 Hz, 2H), 4.55 - 4.98 (m, 2H), 3.07 - 3.11 (m, 2H)

### Chromatograms and NMR spectra

All characterization data are taken from reports or notebook entries provided by WuXi AppTec.

*Chiral HPLC analysis of **2***

Instrument: Waters Arc; column: CHIROBIOTIC T2 (250 x 4.6 mm, 3 µm); mobile phase: A water and B MeOH; gradient: B% = 20% for 8 min; flow rate: 0.6 mL/min; column temperature: 30 °C

Peak 1 fractional area (4.495 min): 99.64%; peak 2 fractional area (5.202 min): 0.36%

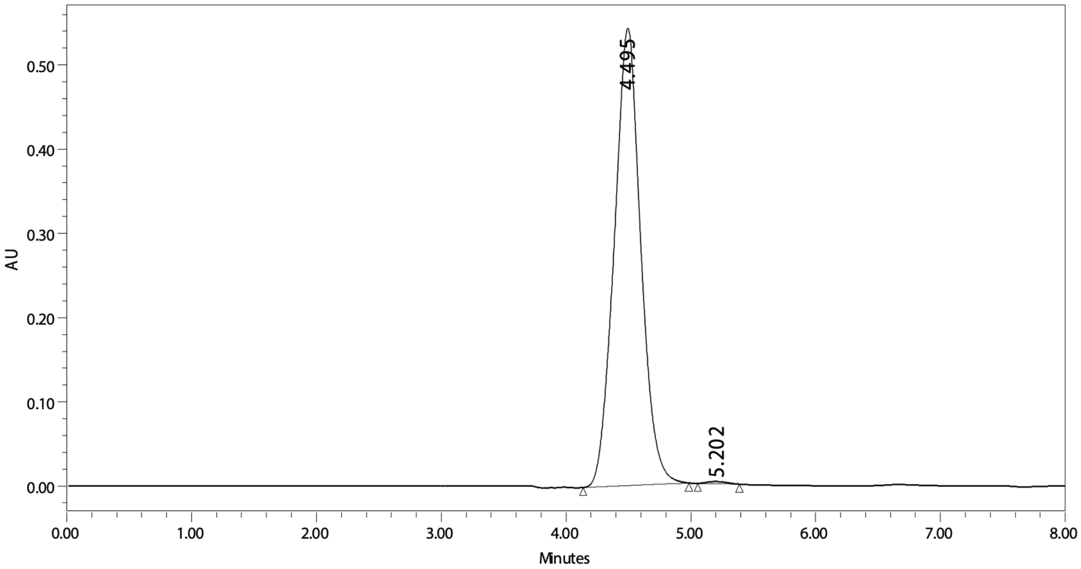


*$^{1}H$ NMR spectrum of **2** (400 MHz, DMSO-$d_6$)*

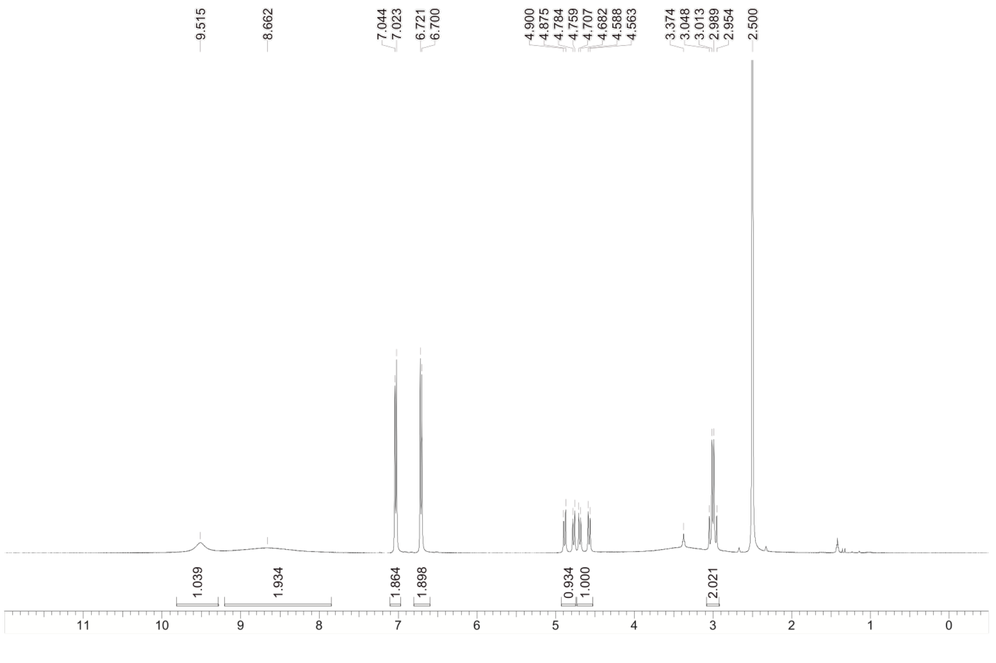

*Chiral HPLC analysis of **3***

Instrument: Waters Arc; column: CHIROBIOTIC T (250 x 4.6 mm, 3 µm); mobile phase: A water and B MeOH; gradient: B% = 50% for 6 min; flow rate: 0.8 mL/min; column temperature: 30 °C

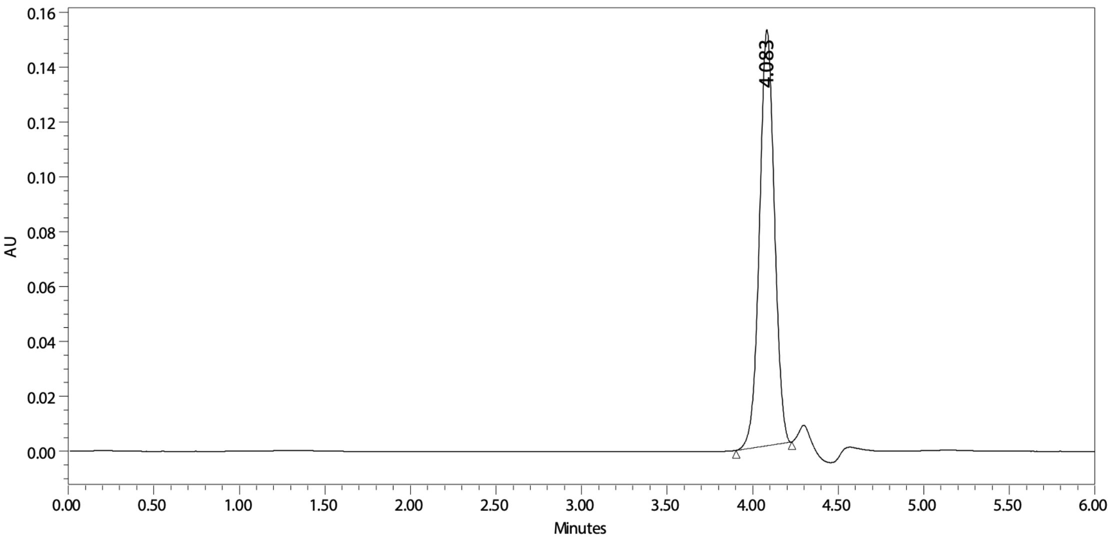


*$^1$H NMR spectrum of **3** (400 MHz, DMSO-$d_6$)*

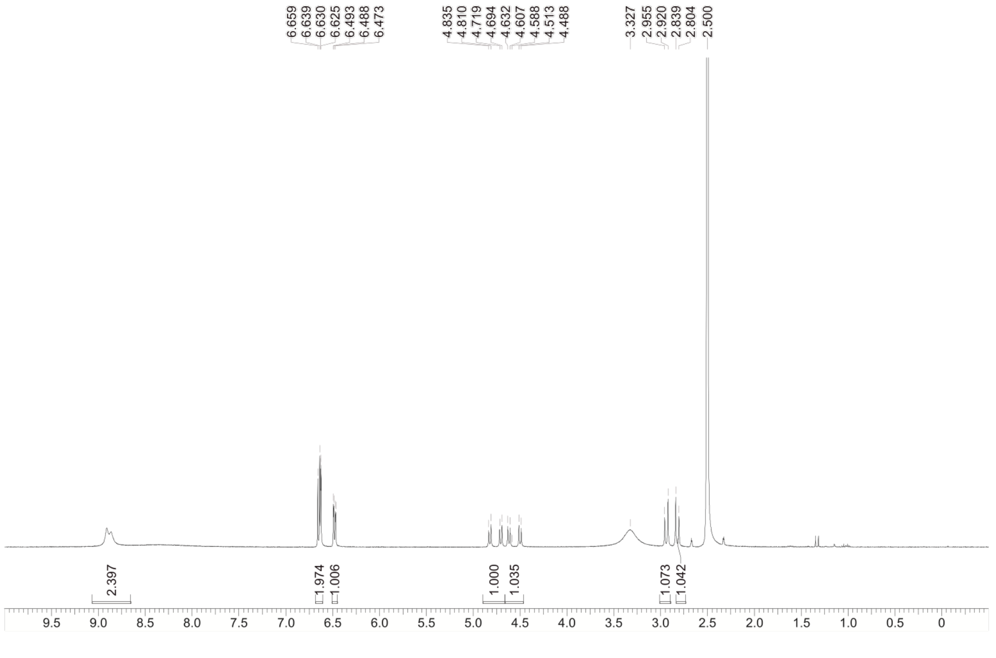

*Chiral HPLC analysis of **26***

Instrument: Waters Arc; column: CHIROBIOTIC T2 (250 x 4.6 mm, 3 µm); mobile phase: A water and B MeOH; gradient: B% = 50% for 15 min; flow rate: 0.8 mL/min; column temperature: 30 °C

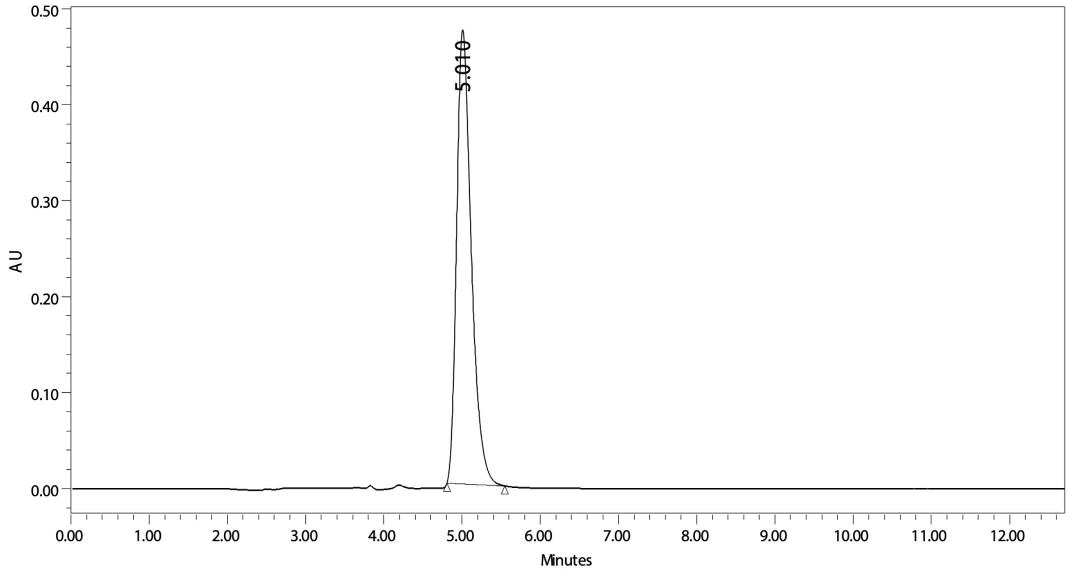


*$^1$H NMR spectrum of **26** (400 MHz, DMSO-$d_6$)*

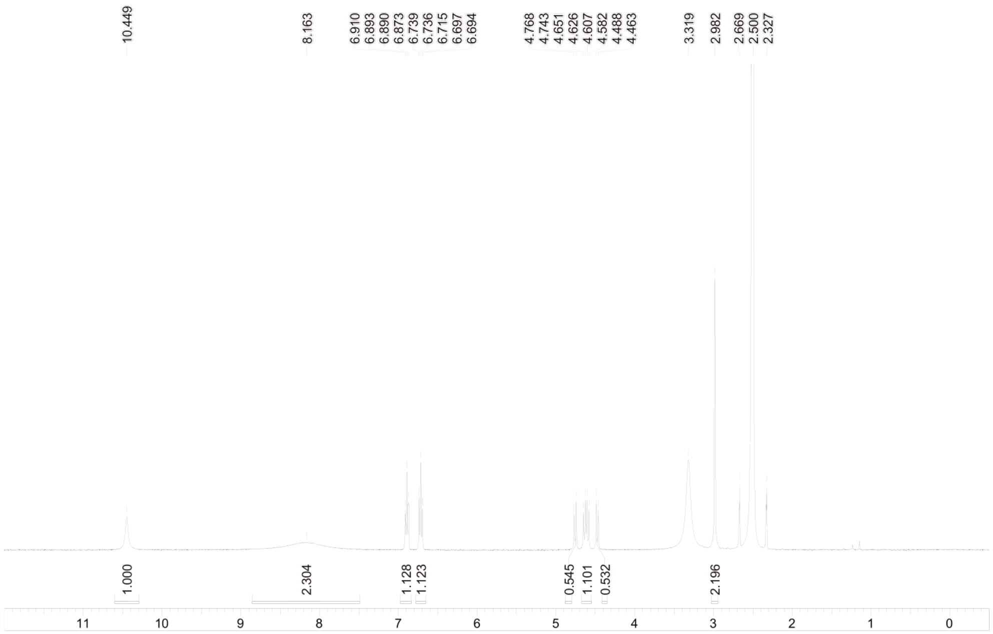

*Chiral HPLC analysis of* ***27***

Instrument: Waters Arc; column: CHIROBIOTIC T2 (250 x 4.6 mm, 3 µm); mobile phase: A water and B MeOH; gradient: B% = 50% for 15 min; flow rate: 0.8 mL/min; column temperature: 30 °C

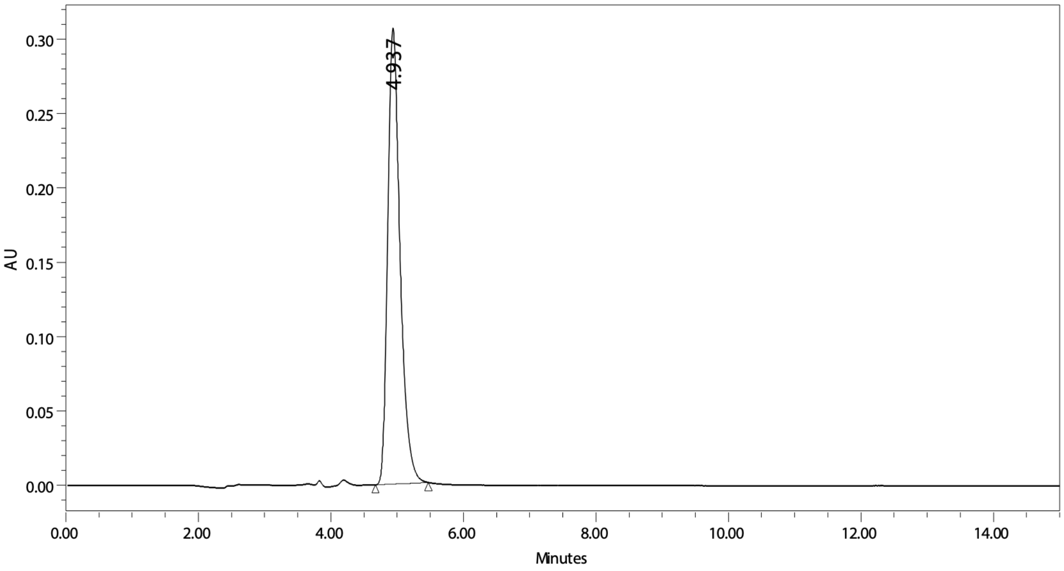


*$^1$H NMR spectrum of* ***27*** *(400 MHz, DMSO-$d_6$)*

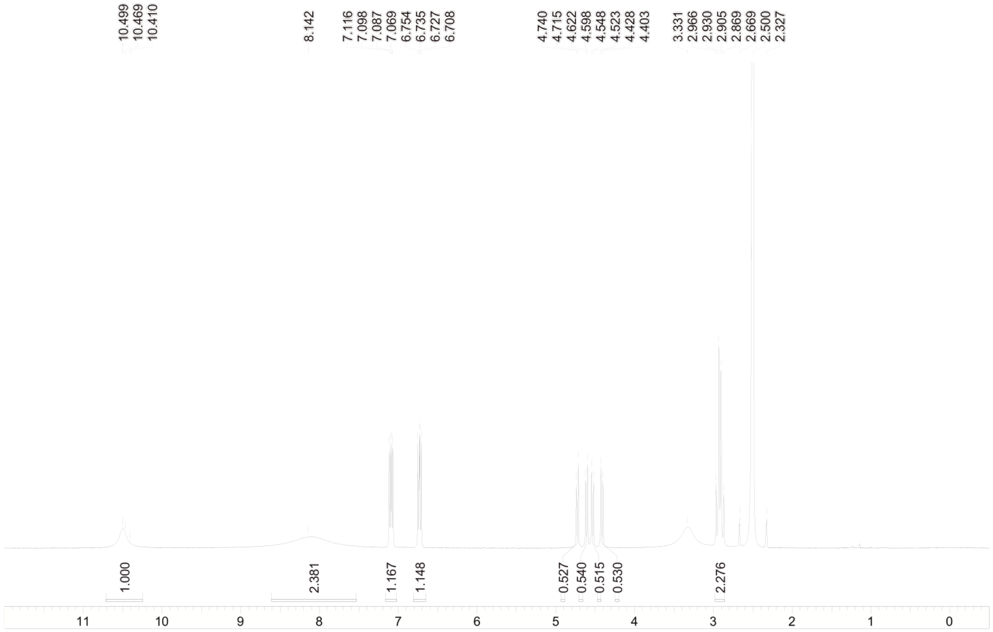

*Chiral SFC analysis of **28***

Instrument: Waters UPC2; column: CHIRALPAK AD-3 (50 x 4.6 mm, 3 µm); mobile phase: A $CO_2$ and B MeOH (0.2% $NH_3$, 7 M in MeOH); gradient: B% = 5% for 0.2 min, 5-50% over 1 min, 50% for 1 min, 50-5% over 0.4 min, 5% for 0.4 min; flow rate: 3.4 mL/min

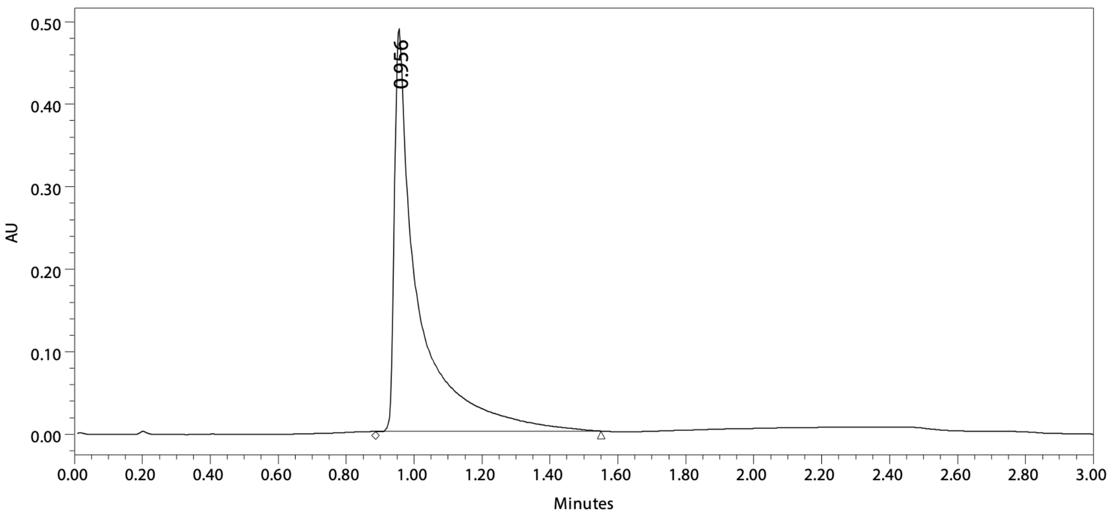


*[1]H NMR spectrum of **28** (400 MHz, DMSO-$d_6$)*

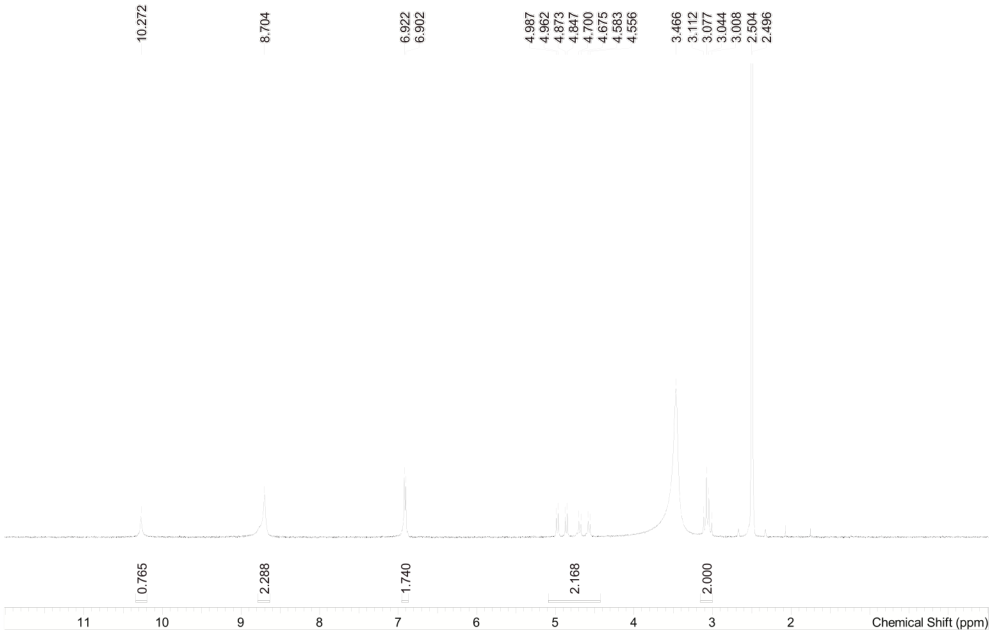
